# Indocyanine green excitation-emission matrix characterization: excitation-dependent emission shifts and application-specific spectra

**DOI:** 10.1101/2025.08.12.667954

**Authors:** Alberto J. Ruiz, Sophie A. Lyon, Ethan P.M. LaRochelle, Kimberley S. Samkoe

## Abstract

**Significance:** Indocyanine green (ICG) is the most widely used fluorophore in fluorescence-guided surgery (FGS), yet its spectral response depends on microenvironment, with implications for system design, inter-system comparisons, and phantom development.

**Aim:** To characterize ICG with excitation–emission matrices (EEMs) in the microenvironments of dimethyl sulfoxide (DMSO), bovine serum albumin (BSA) solutions, and 3D-printed (3DP) resin, and assess excitation-dependent emission, including red-edge excitation shifts (REES) and departures from Kasha’s rule of excitation-independent emission.

**Approach:** EEMs and absorbance spectra were acquired with extracted excitation spectra, emission spectra, emission peaks, centroids, and integrated emission areas under the curve (AUCs). Concentration-dependent behavior was examined in DMSO, and albumin concentration dependence was assessed from 5–100 mg/mL. Data processing employed robust local regression to mitigate excitation scattering artifacts.

**Results:** ICG in DMSO exhibited excitation-independent emission consistent with Kasha–Vavilov behavior. In contrast, ICG in BSA solution and 3DP resin displayed excitation-dependent emission with pronounced REES and additional non-linear departures from Kasha’s rule. To our knowledge, this represents the first documentation of REES and broader anti-Kasha effects for ICG or any FGS fluorophore. Within the excitation range most relevant to ICG-FGS (∼760–805 nm), emission spectra of the BSA solution and 3DP resin overlapped closely, with similar AUC-based comparisons, suggesting that ICG in 3DP resin can serve as a suitable surrogate reference for albumin-bound ICG.

**Conclusions:** The EEM characterization shows that excitation-dependent behavior is a defining feature of ICG in biologically relevant environments, demonstrating that emission cannot be assumed to follow classical Kasha–Vavilov behavior. Reliable comparisons and imaging system design therefore require spectra acquired at defined excitation wavelengths with AUC integration within the emission detection band. Excitation-specific spectra from EEMs establish a consistent framework for inter-system comparisons and phantom standards, while the resulting datasets provide a practical reference for addressing excitation-dependent behavior in ICG sensing applications.

## 1 Introduction

Understanding the spectral characteristics of a fluorophore is crucial for fluorescence sensing,^1^ including fluorescence-guided surgery (FGS).^2^ The microenvironment of a fluorophore can significantly influence its physical and chemical behaviors, with factors such as local viscosity, polarity, temperature, redox conditions, and acidic-base status playing important roles.^3^ In fluorescence imaging, the most relevant changes associated with the fluorophore’s microenvironment are the excitation-emission spectral shifts and variations in the quantum yield,^4^ since they affect the sensitivity and specificity of fluorescence sensing devices.^5–8^ Among the microenvironment-sensitive spectral behaviors, solvatochromism reflects solvent-dependent differential stabilization of the ground and excited states and is observed as steady-state shifts in absorption and emission maxima.^9,10^ A complementary dynamic effect is the red-edge excitation shift (REES), in which the emission spectrum becomes excitation-wavelength dependent when solvent or protein relaxation occurs on timescales comparable to or slower than the fluorescence lifetime.^11–13^ This excitation dependence represents a departure from Kasha’s (Kasha-Vavilov) rule,^14,15^ which assumes excitation-independent emission for a given excited electronic state.^16–20^ Additionally, excitation-dependents shift may be especially relevant in the complex environment of biological systems, where proteins such as serum albumin are present.^21^ As FGS continues to develop, comprehensive characterization of how fluorophore spectral properties respond to physiologically relevant microenvironments can help optimize optical system design, support inter-system comparisons, and guide the development of imaging phantoms and targets that support clinical translation.^7,22,23^

Indocyanine green (ICG) is the most widely used fluorophore in FGS, utilized for tissue perfusion, cardiac flow indication, and lymph node mapping.^24–26^ Its prominence in fluorescence guidance stems from its absorption and emission in the near-infrared (NIR) range, low toxicity, and extensive medical use for more than half a century.^24,25,27,28^ Understanding the variability of ICG’s spectral characteristics within *in vivo* environments can help optimize imaging system design and cross-system comparisons. This includes standard solvent effects as well as non-linear shifts, such as REES. Solvatochromism of ICG has been widely studied and reported. ICG shows pronounced solvatochromism: solvent polarity and binding partners modulate its absorption/emission maxima and quantum yield; albumin binding typically red-shifts and stabilizes the spectra, while aqueous self-aggregation blue-shifts absorption and quenches fluorescence.^29–34^ These environment-dependent shifts have been leveraged in various applications, including human serum albumin-ICG complexes to boost tumor-to-background contrast, conjugates for targeted imaging, liposomal and J-aggregate strategies to enhance stability/brightness, and solvatochromic readouts to map solvent composition and microenvironments in situ.^35–44^ Despite this extensive work on ICG solvatochromism, aggregation, and protein binding, we found no peer-reviewed reports documenting REES or other anti-Kasha excitation-dependent effects in ICG; additionally, comprehensive spectroscopy papers and clinical imaging reviews do not describe excitation-dependent emission consistent with REES or with departures from Kasha’s rule.^26,29–31,37,41,45–48^ By contrast, REES is well established in the literature for proteins, membranes, and protein–ligand complexes, where slow environmental relaxation relative to the fluorescence lifetime produces excitation-dependent red-shifts that can resolve microstate heterogeneity.^49–55^ Differences in excitation–emission behavior under varying environments can be captured by acquiring excitation–emission matrices (EEMs),^56^ which provide extended spectral information that support the assessment of solvatochromism, REES, and other microenvironmental effects..^12^

Excitation–emission matrices (EEMs) are three-dimensional datasets that map fluorescence intensity as a function of excitation and emission wavelength. They are typically acquired by recording emission spectra over a series of discrete excitation wavelengths (**Fig 1a,b**). Conceptually, an EEM can be viewed either as a stack of emission spectra at different excitation wavelengths or, equivalently, as a set of excitation spectra at fixed emission wavelengths (**Fig 1c,d**). This broad mapping of the excitation-emission properties provides more comprehensive information than conventional excitation-emission graphs collected at a single probing wavelength, enabling assessment of excitation-dependent phenomena such as REES. For fluorophores exhibiting excitation-dependent spectral shifts, EEMs allow derivation of emission spectra specific to the excitation characteristics of a given sensing system. A significant limitation of EEMs is the long acquisition times associated with obtaining fluorescence spectra across various excitation wavelengths. However, recent advances in CCD-based fluorescence spectrometers have enabled the rapid acquisition of these data sets. These developments provide a promising pathway for utilizing EEMs as a preferred alternative to individual spectra acquisitions, given their ability to characterize broader aspects of fluorophore behavior.

**Fig 1.**
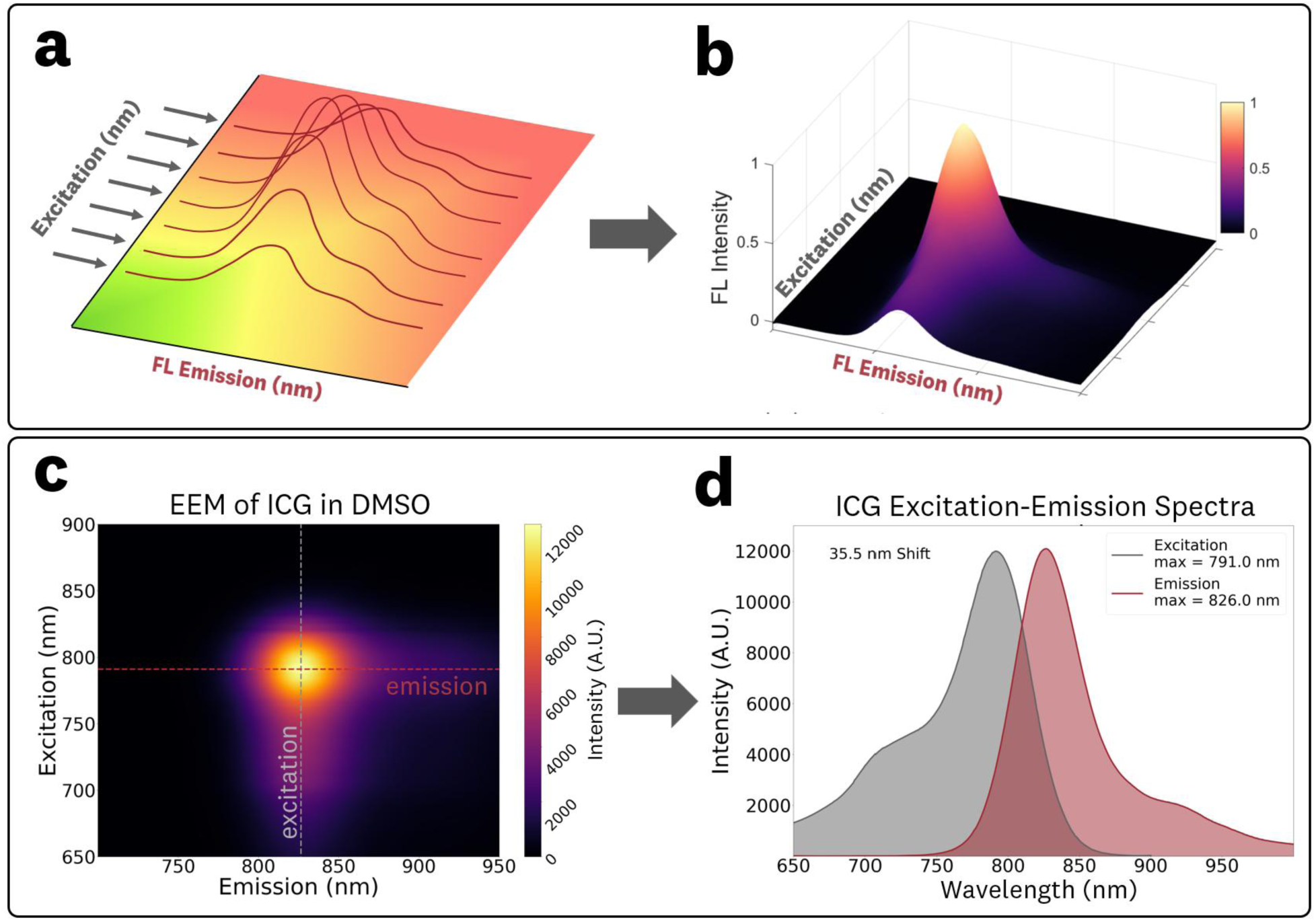
The excitation-emission matrix (EEM) of a fluorophore is generated by **(a)** acquisition of fluorescence emission spectra at various excitation wavelengths to produce the **(b)** three-dimensional EEM data set. The **(c)** EEM can be used to obtain conventional **(d)** excitation and emission spectra by isolating data along a single wavelength on the excitation or emission axis, respectively.

Furthermore, because FGS system performance can affect clinical decision-making,^5,6^ there is a compelling need for imaging standards that can help provide ‘ground truth’ for system characterization.^7,57^ In this context, understanding how the excitation and emission of fluorescence imaging targets can mimic *in vivo* fluorescence is essential to provide relevant one-to-one equivalence to clinical fluorescence imaging applications.

Here, we report excitation–emission matrices of ICG in three microenvironments: dimethyl sulfoxide (DMSO), bovine serum albumin (BSA) solutions, and 3D-printed resin. These measurements establish comprehensive EEM characterization of ICG that reveal red-edge excitation shifts (REES) and departures from Kasha’s rule for ICG in serum albumin and 3D-printed resin, representing, to our knowledge, the first report of this phenomenon for ICG and for any FGS fluorophore. The EEM datasets are provided as supplementary files to support reproducibility, facilitate cross-system benchmarking, and serve as a reference for optical system development. Collectively, the results highlight new considerations for generating excitation-specific spectra, guiding FGS system design and comparison, and informing phantom development for clinically relevant fluorescence imaging.

## 2 Methods

Preparation of the DMSO, BSA, and 3D-printed samples are described in Sections 2.1–2.3. Procedures for the fluorescence EEM acquisition, absorbance measurements, and data processing are provided in Sections 2.4–2.6. A complete list of the tested samples is provided in **Table S1** of the Supplementary Material. The fluorescent samples were prepared using IR-125 laser dye (Exciton Inc., 09030), which is the laser dye marketing name of ICG (sharing the same Chemical Abstracts Service number, 3599-32-4), with identical chemical, quantum yield, and fluorescence spectral characteristics.^58^ IR-125 dye was utilized for this study, rather than clinical ICG dye, due to the availability of purity specifications from the respective manufacturers. Throughout the remainder of this study, the terms IR-125 and ICG are used synonymously.

### 2.1 Preparation of DMSO solution samples

EEM and absorbance measurements for the DMSO solutions were performed on standard 4.5 mL cuvettes (12×12×45 mm, 10 mm pathlength). A 1000 µM ICG stock solution was prepared in DMSO (Sigma-Aldrich, 472301) to pre-suspend the fluorophore and minimize error during spectral comparisons.

For sample preparation, the 1000 µM stock was diluted with DMSO to yield 100 mL of a 1 µM working solution. A total of 3.5 mL of this 1 µM solution was transferred into a cuvette for data collection, with a matched control cuvette containing 3.5 mL of DMSO (0 µM ICG).

To study concentration-dependent EEM effects, an additional dilution of 100 µM was prepared from the 1000 µM DMSO stock. This 100 µM ICG stock was used to create, through serial dilution, 10 mL samples at concentrations of 10, 3, 1, 0.3, 0.1, and 0.03 µM of ICG in DMSO. Volumes of 3.5 mL of each of these solutions were deposited into cuvettes for data collection with an additional control cuvette of 3.5 mL of DMSO (0 µM ICG).

### 2.2 Preparation of BSA solution samples

EEM and absorbance measurements for the BSA solutions were performed on standard 4.5 mL cuvettes (12×12×45 mm, 10 mm pathlength). The 1 µM ICG in BSA solution (44 mg/mL) cuvette sample was prepared using the 1000 µM ICG in DMSO stock (Section 2.1), 150 mg/mL BSA stock, and phosphate-buffered saline (PBS). 10× concentrated PBS (Sigma-Aldrich, P7059) was diluted tenfold with distilled water to prepare 1× PBS. Lyophilized BSA powder (Sigma-Aldrich, A2153) was dissolved in the 1x PBS to the 150 mg/mL BSA stock. From these stocks, 100 mL of 1 µM ICG in BSA solution (44 mg/mL BSA) was prepared; 3.5 mL was transferred to a cuvette for data collection, with a matched control cuvette containing 3.5 mL of BSA solution (44 mg/mL, 0 µM ICG). The ICG in BSA solution was incubated at room temperature and protected from light for an hour prior to measurement. It should be noted that the 1000 µM ICG in DMSO stock solution (Section 2.1) was used in this preparation to mitigate the aggregation of dye in PBS solution, which was experimentally observed at these high concentrations. The 44 mg/mL BSA concentration was used as a representative concentration of albumin in human plasma.^59,60^

To study the effect of serum albumin concentration on spectral properties, 1 µM ICG samples were prepared at BSA concentrations of 5, 10, 25, 50, and 100 mg/mL. First, the 1000 µM ICG in DMSO stock (Section 2.1) was diluted with distilled water to prepare a 9 µM ICG aqueous stock. Separately, BSA was diluted with distilled water to generate two sets of 3.5 mL solutions at each target concentration (5-100 mg/mL): one set served as controls (0 µM ICG), and the other set received 389 µL of the 9 µM ICG stock to yield 3.5 mL cuvette samples at 1 µM ICG. Samples containing ICG were incubated at room temperature, protected from light, for 1 h prior to measurement.

### 2.3 Preparation of 3D-printed samples

The 1 µM ICG in 3D-printed (3DP) resin and the control (0 µM) samples were prepared using a proprietary clear photocurable resin (MML-REPC-001, QUEL Imaging) and the 1000 µM ICG in DMSO stock solution (Section 2.1). The 3DP samples were prepared following a previously published method.^22^ In brief, 100 µL of the ICG stock was mixed into 100 mL of the clear resin and printed using layer-by-layer stereolithography printing (405 nm curing) to produce a fluorescent 3DP cuvette (12×12×40 mm^3^) to match the dimensions of the standard cuvette samples; a control cuvette (0 µM ICG) was also printed using the same method. After printing, the cuvettes were cleaned with isopropyl alcohol and post-cured utilizing a high-radiance 385 nm light (Solarez, 88903). To minimize scattering of light, the 3DP cuvettes were progressively wet-sanded with distilled water and 1500, 2000, and 3000 grit silicon carbide sandpaper, followed by a two-step liquid polisher (Novus 7100). Sanding and polishing resulted in an optically clear surface, enabling data collection comparable to liquid samples prepared in a standard cuvette.

### 2.4 Fluorescence excitation-emission matrix acquisition

Fluorescence EEMs and absorbance spectra were collected using a dual fluorescence and absorbance spectrometer (Duetta, HORIBA Scientific). This spectrometer uses a xenon arc lamp with a monochromator, enabling excitation and absorbance sweeps over the 250–1000 nm range at 1 nm steps. It is equipped with a linear CCD sensor for fluorescence emission capture (250–1050 nm) and a silicon photodiode for absorbance measurements (250 – 1000 nm), utilizing right-angle (90°) excitation-emission geometry. For EEM acquisition, samples were excited from 650 to 900 nm in 1 nm increments, and the resulting fluorescence emission was collected over 600–1000 nm with a spectral resolution of ∼0.5 nm (1 pixel). All spectra were collected using a 3 nm excitation and emission bandpass slit size and three measurement accumulations. The reported maximum fluorescence emission values were assumed to have an uncertainty of ±1 nm, consistent with the manufacturer’s specified wavelength accuracy. Although the CCD provided a sampling resolution of ∼0.5 nm per pixel, the absolute calibration accuracy was ±1 nm and therefore taken as the limiting factor. Excitation values were likewise assumed to have ±1 nm uncertainty, based on acquisition step size and manufacturer specifications. The uncertainty of the reported Stokes shifts was assumed to be ±1.4 nm, obtained by adding the excitation and emission uncertainties in quadrature. Integration times of 1.0–2.0 s were used, with identical integration times maintained for comparative measurements to ensure consistent cross-sample fluorescence emission comparisons. For the varying concentration measurements of DMSO solutions (Section 2.1), the 3 µM and 10 µM samples were recorded with half the integration time to avoid detector saturation; their measured intensity counts were subsequently doubled during data processing to account for integration time differences.

The reported EEMs are calculated by subtracting the acquired data of the corresponding control (0 µM ICG) sample from the fluorescent sample. The emission spectra at a given excitation were obtained from plotting the intensity values along the emission axis of the EEM (**Fig 1c,d**). The excitation spectra at a “monitored” fluorescence wavelength were derived from plotting the intensity values along the excitation axis of the EEM (**Fig 1c,d**). The integrated fluorescence emission intensity at a given excitation wavelength, referred to as areas under the curve (AUCs) for the remainder of the paper, were calculated by trapezoidal numerical integration (implemented using the *trapz* function from the NumPy library) of extracted emission spectra (650–1000 nm), with emission centroids defined as the equal-area wavelength corresponding to the integrated distribution. Emission peak maxima were determined as the highest intensity values of the processed spectra (see Section 2.6 for details on data processing).

### 2.5 Absorbance spectra acquisition

Absorbance spectra were collected over the 600-1000 nm range at 1 nm step increments utilizing the spectrometer silicon photodiode detector (see Section 2.4 for instrument details). For all samples, an integration time of 0.1 s and a band pass slit size of 3 nm were used with corresponding blanks of DMSO, BSA, and 3DP resin. The reported max absorbance values are assumed to have a ± 1 nm uncertainty, given the corresponding acquisition step size parameters and manufacturers wavelength accuracy specification. Absorbance data is reported with units of optical density (OD), where the measured transmission = 10^-OD^. The manufacturer provides an absorbance accuracy specification of ± 0.02 OD.

### 2.6 Data processing

The EEM and absorbance spectra were processed using robust local regression smoothing with a span of 40 nm. The particular method utilized, robust locally estimated scatterplot smoothing (RLOESS), uses robust locally weighted linear least squares regression with a 2^nd^ degree polynomial model.^61,62^ This robust smoothing allows for the elimination of outlier measurements and generally provided better peak estimations than traditional smoothing techniques (i.e. Savitzky-Golay filtering) when tested against Gaussian and skewed Gaussian fits to the spectral data. Furthermore, this robust smoothing provided the ability to eliminate scattering peaks introduced by the BSA and 3D printed resin. An example of this scattering artifact elimination by the robust smoothing is shown in **Fig 2**. RLOESS can be implemented through the use the *smooth* function in MATLAB or, equivalently, in Python with a modified implementation of the Pypi.org *loess* project,^63,64^ which provides equivalent robust weights to the native MATLAB implementation.

**Fig 2.**
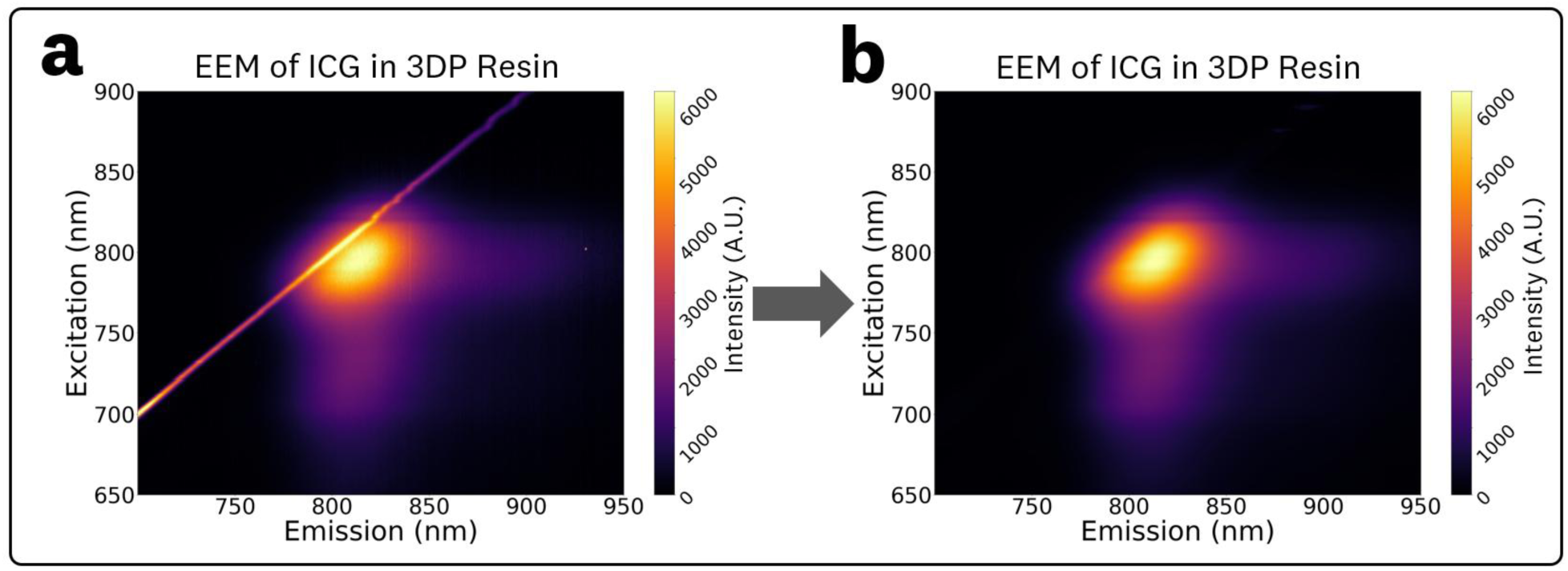
EEMs for samples with inherent scattering (3DP resin and albumin solutions) exhibit a scattering peak along the excitation-emission line **(a)** which can be corrected through the use of the RLOESS smoothing algorithm to result in the post-processed EEMs **(b)**.

## 3 Results

### 3.1 ICG in DMSO

The acquired EEM for 1 µM ICG in DMSO and the associated spectra are shown in **Fig 3**. The EEM (**Fig 3a**) maxima for excitation and emission were measured as 792 nm and 826.5 nm, respectively. The corresponding excitation and emission spectra at these maxima are plotted in **Fig 3b**, showing a measured Stokes shift of 34.5 nm.

**Fig. 3.**
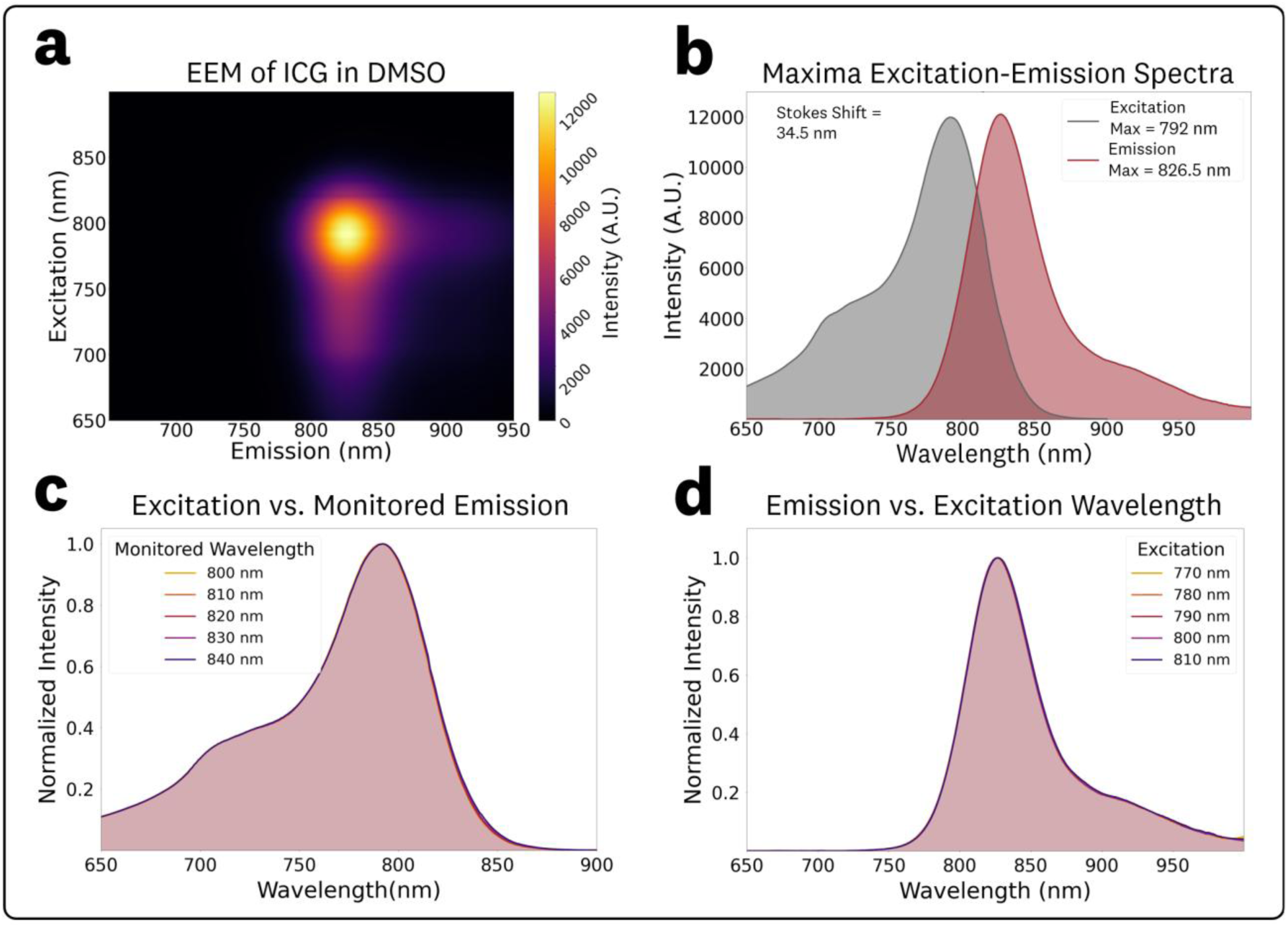
EEM and associated spectra for 1 µM ICG in DMSO: **(a)** Top-down plot of the acquired EEM. **(b)** Excitation and emission spectra extracted from the EEM maxima. **(c)** Normalized excitation spectra at varying monitoring emission wavelengths in the 800–840 nm range at 10 nm steps and **(d)** Normalized emission spectra at varying excitation wavelengths in the 770–810 nm range at 10 nm steps.

To assess potential spectral shifts in the excitation, spectra corresponding to monitored emission wavelengths between 800 and 840 nm (10 nm steps) were extracted from the EEM data (**Fig 3c**). The identical spectral shapes of the excitation spectra indicate there are no significant shifts in the fluorescence spectral characteristics for varying monitored emission wavelengths.

To evaluate excitation-dependent, spectra from excitation wavelengths in the 770–810 nm range (10 nm steps) were extracted from the EEM data (**Fig 3d**). The identical spectral shapes of the fluorescence emission indicate there are no significant shifts in the fluorescence spectral characteristics for varying excitation wavelengths. The measured emissions maxima, relative peak intensity, centroid, and relative integrated fluorescence AUC for the various excitations are provided in **Table 1**. No excitation-dependent shifts were observed in either the emission peak or centroid, confirming that ICG in DMSO follows classical Kasha–Vavilov behavior.

**Table 1:** Emission maxima, relative peak intensities, emission centroids, and relative integrated emission AUC extracted from the EEM of 1 µM ICG in DMSO at excitation wavelengths of 770-810 nm (10 nm steps).

| Excitation (nm) | Emission Peak | Peak Intensity | Emission Centroid | AUC |
| --- | --- | --- | --- | --- |
| 770 nm | 826.5 nm | 0.73 | 836.5 nm | 0.73 |
| 780 nm | 826.5 nm | 0.91 | 836.5 nm | 0.91 |
| 790 nm | 826.5 nm | 1.00 | 836.5 nm | 1.00 |
| 800 nm | 826.5 nm | 0.95 | 836.5 nm | 0.95 |
| 810 nm | 827.0 nm | 0.74 | 837.0 nm | 0.75 |

The acquired EEMs of ICG in DMSO at varying concentrations of 0.03, 0.1, 0.3, 1.0, 3.0, and 10 µM are provided in **Figure S1** of the Supplementary Material. These measurements showed fluorescence quenching at the 10 µM concentration (**Fig 4a**), concentration-dependent red shifts (CDRS) (**Fig 4b**), and disparities between absorbance and emission spectra due to inner filter effects (IFE) (**Fig 4c,d**). The log-log plot of maximum fluorescence intensity vs. concentration (**Fig 4a**) shows a constant increase in fluorescence emission vs. concentration for the 0.03–3 µM range with a drop in intensity for the 10 µM concentration. This drop in fluorescence intensity at the highest concentration is most likely attributed to both primary and secondary inner filter effects (IFE), which correspond to quenching caused by attenuation of the excitation beam and re-absorption of emitted fluorescence, respectively.^65^

**Fig. 4:**
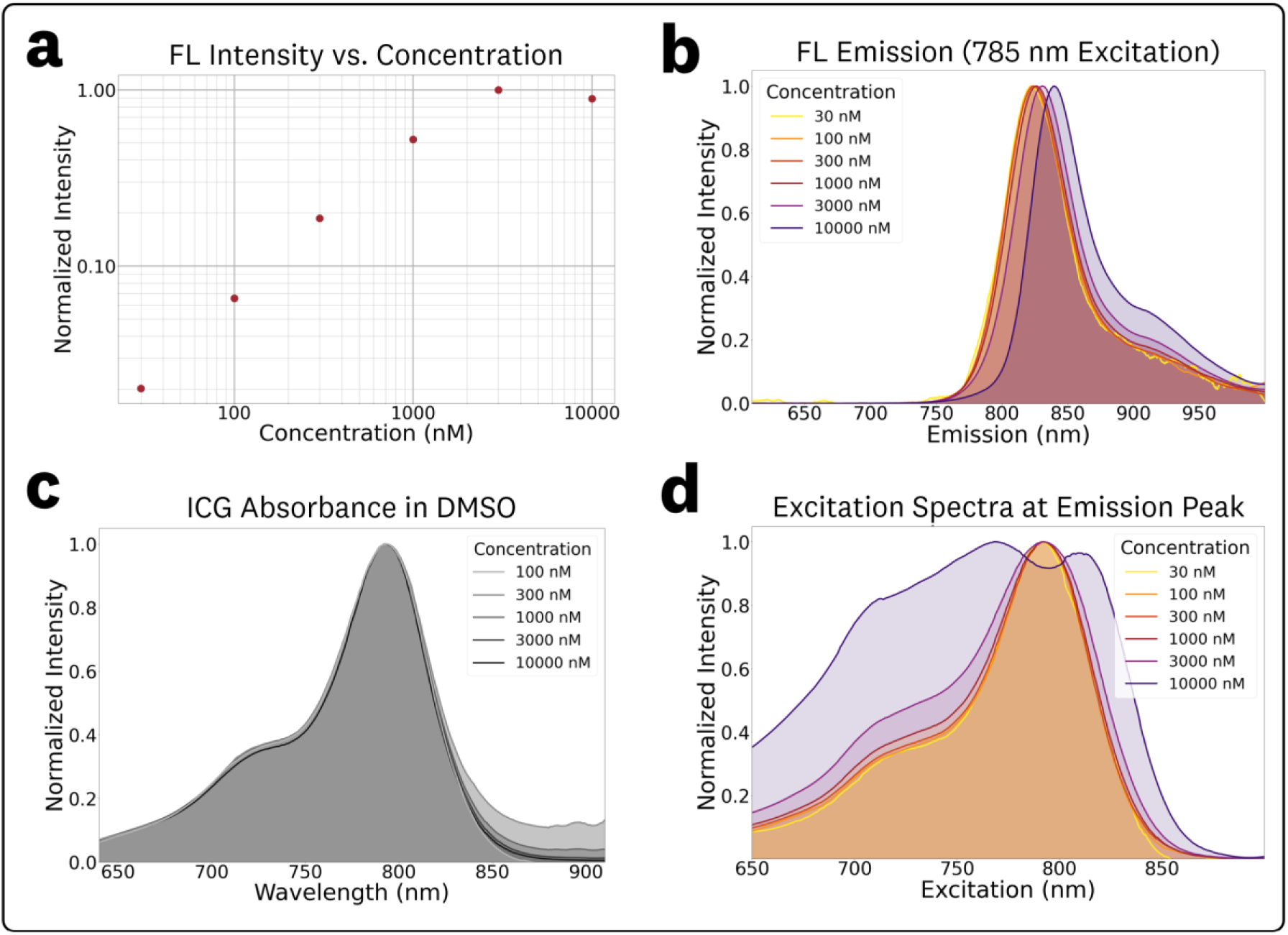
Concentration-dependent spectral effects for ICG in DMSO. **(a)** Plot of maximum fluorescence intensity vs. concentration showing a consistent increase for the 0.03–3 µM range with a drop-off in intensity for the 10 µM concentration. **(b)** Normalized fluorescence emission spectra at 785 nm excitation showing concentration-dependent red shifts. **(c)** Absorbance spectra show consistent spectral features over the entire concentration while the **(d)** excitation spectra show broadening and changes due to fluorescence re-absorption for the 3 µM and 10 µM concentrations.

To simplify discussion of concentration dependent fluorescence shifts, normalized fluorescence emission spectra at 785 nm excitation were extracted from the EEMs (**Fig 4b**). The corresponding measured maximum emission wavelengths and normalized intensities for these plots were: (0.03 µM, 822.0 nm, 0.020), (0.1 µM, 823.0 nm, 0.066), (0.3 µM, 824.0 nm, 0.19), (1 µM, 826.0 nm, 0.52), (3 µM, 830.5 nm, 1.0), and (10 µM, 839.5 nm, 0.89). This red-shift in fluorescence emission maxima for increasing concentrations (termed CDRS), primarily results from the re-absorption of emitted fluorescence.^65^

The measured absorbance data is plotted in **Fig 4c**, showing equivalent spectra from the full 0.03–10 µM concentration range. Data from the 0.03 µM sample was excluded from **Fig 4c** due to the low signal-to-noise ratio of its absorbance acquisition. **Table S2** in the Supplementary Material contains summarized OD absorbance measurements for the varying concentrations, showing no significant variation in calculated molar extinction coefficients (**Table S3** in the Supplementary Material) for the 0.1–10 µM range. In contrast to absorbance, the excitation spectra (**Fig 4d**) at varying concentrations, generated from EEMs at the **Fig 4b** emission peaks, shows broadening and quenching for the 3 µM and 10 µM concentrations. This discrepancy between the absorbance and excitation spectra is caused by secondary IFE effects from the re-absorption of fluorescence emission at these high concentrations.^65^

### 3.2 ICG in BSA Solution

The acquired EEM for 1 µM ICG in BSA solution (44 mg/mL) and associated spectra are shown in **Fig 5**. Compared to the ICG in DMSO EEM (**Fig 3a**), the ICG in BSA solution EEM (**Fig 5a**) showed a significant ‘rotation’ of the central spectra feature, indicating an excitation-dependent REES and, consequently, a departure from Kasha’s rule, with substantial shifts in the emission spectral characteristics across excitation wavelengths. The EEM maxima for excitation and emission were measured as 794.0 nm and 813.5 nm, respectively. The corresponding excitation and emission spectra at these maxima are plotted in **Fig 5b**, showing a measured Stokes shift of 19.5 nm.

**Fig. 5.**
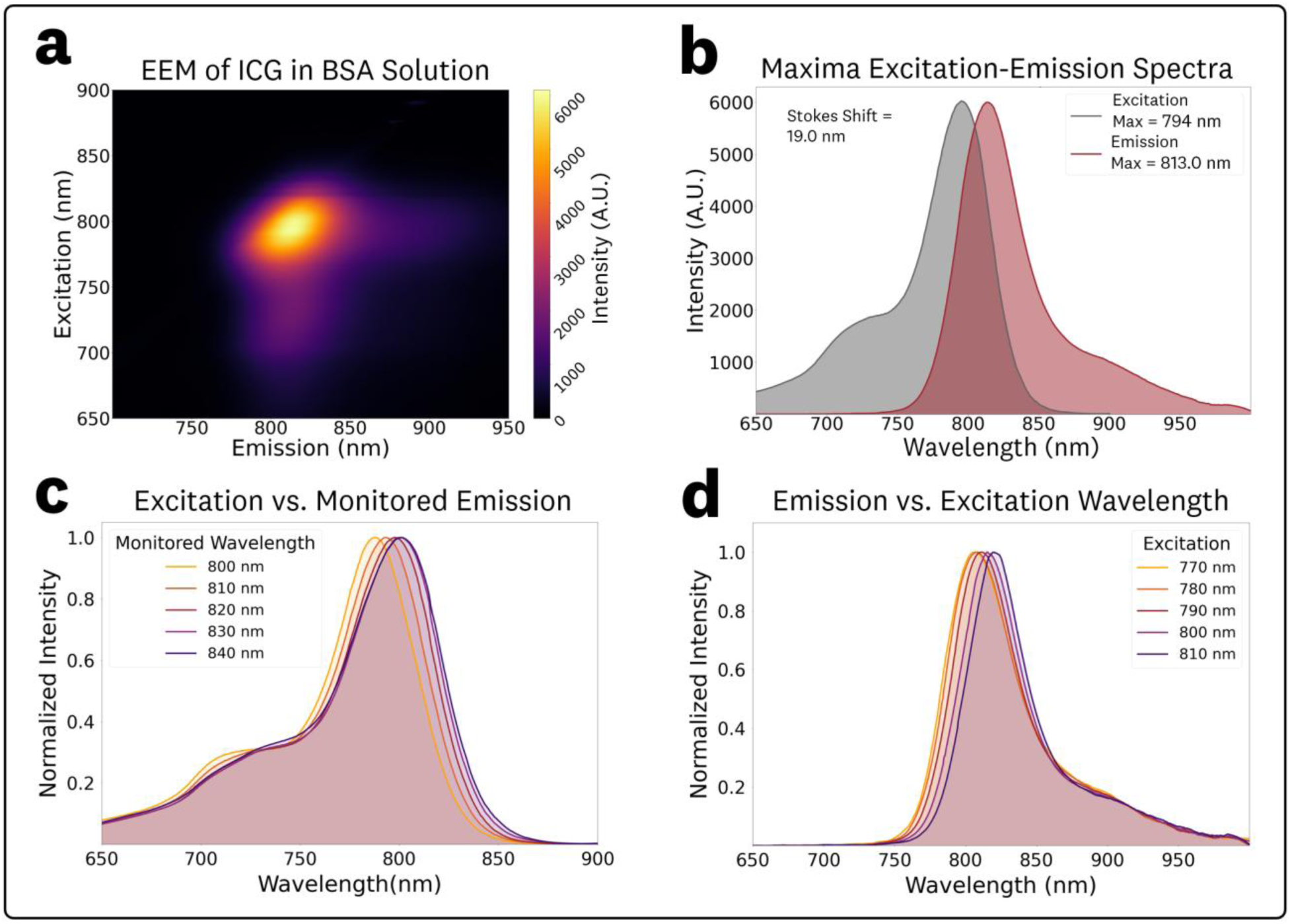
EEM and associated spectra for 1 µM ICG in BSA solution (44 mg/mL). **(a)** Top-down plot of the acquired EEM showing a ‘rotation’ of the central spectral feature indicating excitation-dependent spectral shifts. **(b)** Excitation and emission spectra extracted from the EEM maxima. **(c)** Normalized excitation spectra at varying monitoring emission wavelengths in the 800–840 nm range at 10 nm steps and **(d)** Normalized emission spectra at varying excitation wavelengths in the 770–810 nm range at 10 nm steps.

To assess spectral shifts in excitation, spectra corresponding to monitored emission wavelengths between 800 and 840 nm (10 nm steps) were extracted from the EEM data (**Fig 5c**). The resulting spectra showed increasing shifts in excitation maxima, with measured peaks at 788, 793, 798, 801 and 801 nm for monitored emission wavelengths of 800, 810, 820, 830, and 840 nm, respectively.

To quantify spectral shifts in the fluorescence emission and intensity, spectra from excitation wavelengths in the 760–820 nm range (10 nm steps) were extracted from the EEM data (**Fig 5d**). The measured emissions maxima, relative peak intensities, centroids, and relative AUC values for the various excitations are summarized in **Table 2**. The resulting fluorescence emission spectra showed excitation-dependent changes in the peak wavelength and centroid, with only minor changes in photon distribution (spectral broadening) as indicated by the close agreement between peak intensity and AUC ratios. Over the 770–810 nm excitation range, the emission peak and centroid exhibited average red-shifts of 0.35 and 0.26 nm per nm of excitation, respectively.

**Table 2:** Emission maxima, relative peak intensities, emission centroids, and relative integrated emission AUC extracted from the EEM of 1 µM ICG in BSA solution (44 mg/mL) at excitation wavelengths of 770–810 nm (10 nm steps).

| Excitation (nm) | Emission Peak | Peak Intensity | Emission Centroid | AUC |
| --- | --- | --- | --- | --- |
| 770 nm | 806.5 | 0.62 | 819.5 nm | 0.66 |
| 780 nm | 808.0 | 0.84 | 820.0 nm | 0.88 |
| 790 nm | 811.0 | 0.99 | 822.5 nm | 1 |
| 800 nm | 815.5 | 1.00 | 826.0 nm | 0.96 |
| 810 nm | 820.5 | 0.82 | 829.5 nm | 0.76 |

The acquired ICG EEMs at different concentrations of BSA in the 5–100 mg/mL range are provided in **Figure S2** of the Supplementary Material. The EEM measurements showed no significant changes in spectral characteristics for varying BSA mg/mL concentrations within the tested range, which covers the full range of biologically relevant concentrations.^59,60^ To simplify discussion of BSA concentration dependent fluorescence shifts, normalized fluorescence emission spectra at 785 nm excitation were extracted from the EEMs (**Fig 6**). The excitation plots (**Fig 6a**), obtained by monitoring fluorescence at the emission maximum and normalized by the set maxima, show no significant spectral changes, with all excitation peak maxima at 803 nm. Similarly, the fluorescence emission plots (**Fig 6b**), normalized by the set maxima, show no significant changes in the photon distributions of the emission. The measured maximum emissions and normalized intensities for the various concentrations of BSA were: (5 mg/mL, 816.0 nm, 0.96), (10 mg/mL, 815.5 nm, 0.96), (25 mg/mL, 815.5 nm, 0.97), (50 mg/mL, 815.0 nm, 0.94), and (100 mg/mL, 815.5 nm, 1.0).

**Fig. 6.**
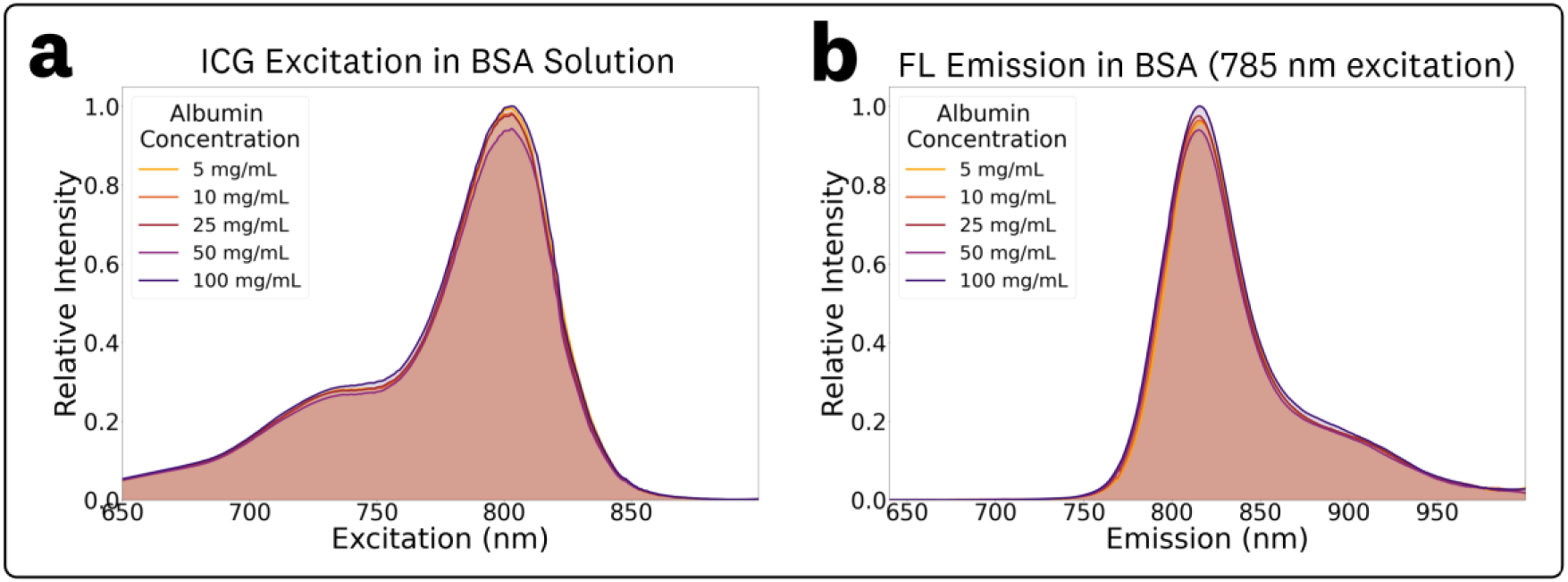
Fluorescence measurements of ICG in 5–100 mg/mL BSA solutions at 785 nm excitation: **(a)** relative excitation spectra obtained at the monitored peak emission wavelength, and **(b)** relative emission spectra, both indicating no significant changes in spectral characteristics across concentrations.

### 3.3 ICG in 3DP Resin

The acquired EEM for 1 µM ICG in 3DP resin and associated spectra are shown in **Fig 7**. Compared to the ICG in DMSO EEM (**Fig 3a**), the ICG in 3DP resin EEM (**Fig 7a**) showed a significant ‘rotation’ of the central spectra feature, indicating an excitation-dependent REES and, consequently, a departure from Kasha’s rule, with substantial shifts in the emission spectral characteristics across excitation wavelengths.^12^ The EEM maxima for excitation and emission were measured as 809 nm and 823.5 nm, respectively. The corresponding excitation and emission spectra at these maxima are plotted in **Fig 7b**, showing a measured Stokes shift of 14.5 nm.

**Fig. 7.**
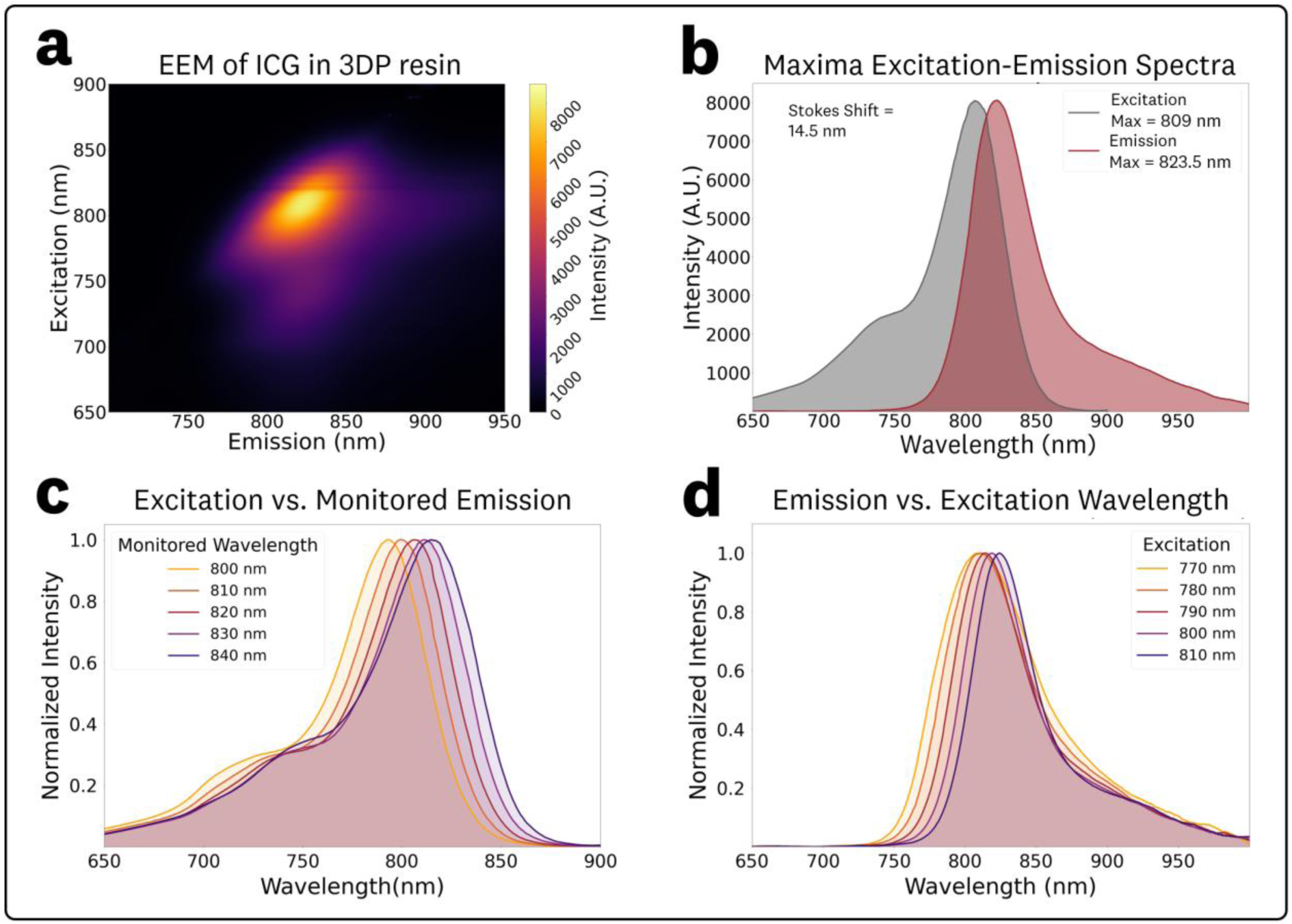
EEM and associated spectra for 1 µM ICG in 3DP resin. **(a)** Top-down plot of the acquired EEM showing a ‘rotation’ of the central spectral feature indicating excitation-dependent spectral shifts. **(b)** Excitation and emission spectra extracted from the EEM maxima. **(c)** Normalized excitation spectra at varying monitoring emission wavelengths in the 800–840 nm range at 10 nm steps and **(d)** Normalized emission spectra at varying excitation wavelengths in the 770–810 nm range at 10 nm steps.

To assess spectral shifts in excitation, spectra corresponding to monitored emission wavelengths between 800 and 840 nm (10 nm steps) were extracted from the EEM data (**Fig 5c**). The resulting spectra showed increasing shifts in excitation maxima, with measured peaks at 793, 799, 806, 811 and 815 nm for monitored emission wavelengths of 800, 810, 820, 830, and 840 nm, respectively.

To quantify spectral shifts in the fluorescence emission, spectra from excitation wavelengths in the 760– 820 nm range (10 nm steps) were extracted from the EEM data (**Fig 7d**). The measured emissions maxima, relative peak intensities, centroids, and relative AUC values for the various excitations are summarized in **Table 3**. The resulting fluorescence emission spectra showed excitation-dependent changes in the peak wavelength and centroid, with noticeable changes in photon distribution (spectral broadening), as reflected by differences between peak intensity and AUC ratios. Over the 770–810 nm excitation range, the emission peak and centroid exhibited average red-shifts of 0.39 and 0.26 nm per nm of excitation, respectively.

**Table 3:** Emission maxima, relative peak intensities, emission centroids, and relative integrated emission AUC extracted from the EEM of 1 µM ICG in 3DP Resin at excitation wavelengths of 770-810 nm (10 nm steps).

| Excitation (nm) | Emission Peak | Peak Intensity | Emission Centroid | AUC |
| --- | --- | --- | --- | --- |
| 770 nm | 809.0 nm | 0.44 | 824.5 nm | 0.62 |
| 780 nm | 810.5 nm | 0.61 | 825.0 nm | 0.76 |
| 790 nm | 815.0 nm | 0.79 | 827.5 nm | 0.9 |
| 800 nm | 819.0 nm | 0.96 | 831.0 nm | 1 |
| 810 nm | 824.0 nm | 1.00 | 835.5 nm | 0.99 |

### 3.4 ICG Spectral Comparison

The absorbance spectra, excitation spectra for the integrated emission AUC, and emission spectra at 760, 785 nm and 805 nm excitation for 1 µM ICG in DMSO, BSA solution (44 mg/mL), and 3DP resin are shown in **Fig 8**, with summarized data provided in **Table 4**. Because the BSA solution and 3DP resin exhibited excitation-dependent emission behavior (**Fig. 5** & **Fig. 7**), excitation-specific spectra provide an appropriate basis for comparing fluorescence responses across microenvironments in fluorescence sensing applications. Accordingly, fluorescence emission spectra for 760, 785, and 805 nm excitation are shown in **Fig. 8c–h**, representing three of the most common excitation wavelengths used for ICG imaging in FGS devices.^7^

**Fig. 8.**
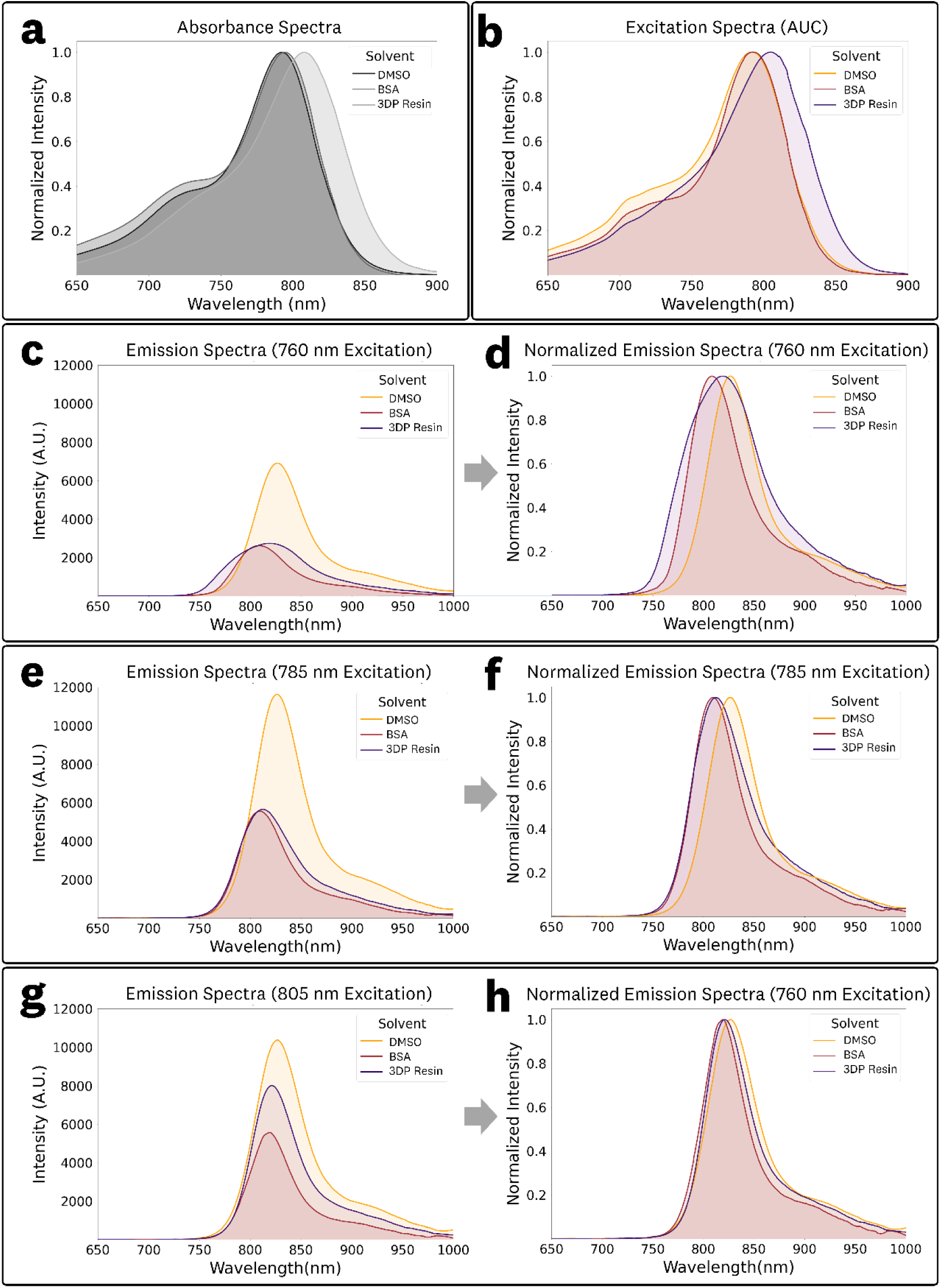
Spectral comparisons of 1 µM ICG in DMSO, BSA solution (44 mg/mL), and 3DP Resin: **(a)** Absorbance spectra, (b) excitation spectra calculated from the EEM emission AUC, **(c,d)** fluorescence emission spectra at 760 nm excitation, **(e,f)** fluorescence emission spectra at 785 nm excitation, and **(g,h)** fluorescence emission spectra at 805 nm excitation

**Table 4:** Summarized data for the comparison of the spectral behavior of ICG in DMSO, BSA solution and 3DP resin microenvironments. The integrated emission AUC is normalized to the counts of the DMSO solution emission maxima at the 785 nm excitation.

|  | Absorbance |  | Excitation | Emission @<br>760 nm excitation |  |  | Emission @<br>785 nm excitation |  |  | Emission @<br>805 nm excitation |  |  |
| --- | --- | --- | --- | --- | --- | --- | --- | --- | --- | --- | --- | --- |
|  | Peak (nm) | OD | Peak (nm) | Peak (nm) | Centroid (nm) | AUC (Norm) | Peak (nm) | Centroid (nm) | AUC (Norm) | Peak (nm) | Centroid (nm) | AUC (Norm) |
| DMSO | 792 | 0.167 | 791 | 826.5 | 836.5 | 0.60 | 826.5 | 836.5 | 1.00 | 827.0 | 837.0 | 0.89 |
| Albumin | 795 | 0.142 | 792 | 808.0 | 821.0 | 0.24 | 810.0 | 821.0 | 0.47 | 819.5 | 828.0 | 0.43 |
| 3DP Resin | 807 | 0.159 | 805 | 819.0 | 826.5 | 0.33 | 813.0 | 826.0 | 0.54 | 821.0 | 833.0 | 0.66 |

The absorbance spectra (**Fig 8a**) showed maxima at 792, 795, and 807 nm for DMSO, BSA solution, and 3DP resin, respectively. The corresponding peak ODs differed by less than 5% in transmission, calculated as 10^-OD^ (**Table 1**). The excitation spectra (**Fig 8b**), calculated from the integrated emission AUC, showed similar differences in peak positions, with maxima at 791, 792, and 805 nm.

The emission spectra demonstrated notable differences in spectral overlap arising from the excitation-dependent behavior of the BSA solution and 3DP resin samples (**Fig. 8d,f,h**). At 760 nm excitation (**Fig. 8c,d**), the 3DP resin spectra was broad, overlapping with both the DMSO and BSA solution, while DMSO and BSA spectra showed limited overlap (**Fig. 8d**). At 785 nm excitation (**Fig. 8e,f**), the spectra of the BSA solution and 3DP showed substantial overlap (**Fig.8f**) with nearly identical AUC intensities (**Fig 8e**, **Table 4**). At 805 nm excitation (**Fig. 8g,h**), the emission spectra of all three samples showed substantial overlap (**Fig. 8h**). Furthermore, the excitation-dependent peak maxima, centroids, and integrated emission AUC, and photon distributions highlight the need to consider the excitation wavelength and emission collection of the fluorescence sensing device to provide relevant comparisons within varying fluorophore microenvironments and system design parameters. The observed excitation-dependent emissions, leading to non-linear shifts in the emission peaks and centroids, are further examined in Section 3.5.

### 3.5 Excitation-Dependency and Red-edge Shift Comparisons

To further explore excitation-dependent effects on the emission spectra, plots of emission peak vs. excitation and emission centroid vs. excitation were generated from the respective EEMs for 1 µM ICG in DMSO, BSA solution (44 mg/mL), and 3DP resin (**Fig. 9**). It is worth noting that the centroid plots showed reduced noise compared to the peak plots due to the variance-reducing effect of AUC integration. For ICG in DMSO (**Fig. 9a,d**), neither the emission peak nor the centroid wavelength exhibited significant variation with excitation wavelength, consistent with classical Kasha rule behavior and excitation-independent emission. In contrast, ICG in BSA solution (**Fig. 9b,e**) and in 3DP resin (**Fig. 9c,f**) showed pronounced shifts in both peak and centroid emission wavelengths as a function of excitation, demonstrating strong non-linear excitation-dependence and red-edge effects. Both the BSA solution and 3DP resin show a local maxima at ∼740-750 nm range, followed by a pronounced REES trend at excitation wavelengths >770 nm. Notably, this localized increase extends beyond the traditional REES behavior, providing strong evidence of anti-Kasha emission dynamics and suggesting that ICG may undergo multiple relaxation pathways depending on excitation wavelength for the BSA solution and 3DP resin microenvironments.

**Fig 9.**
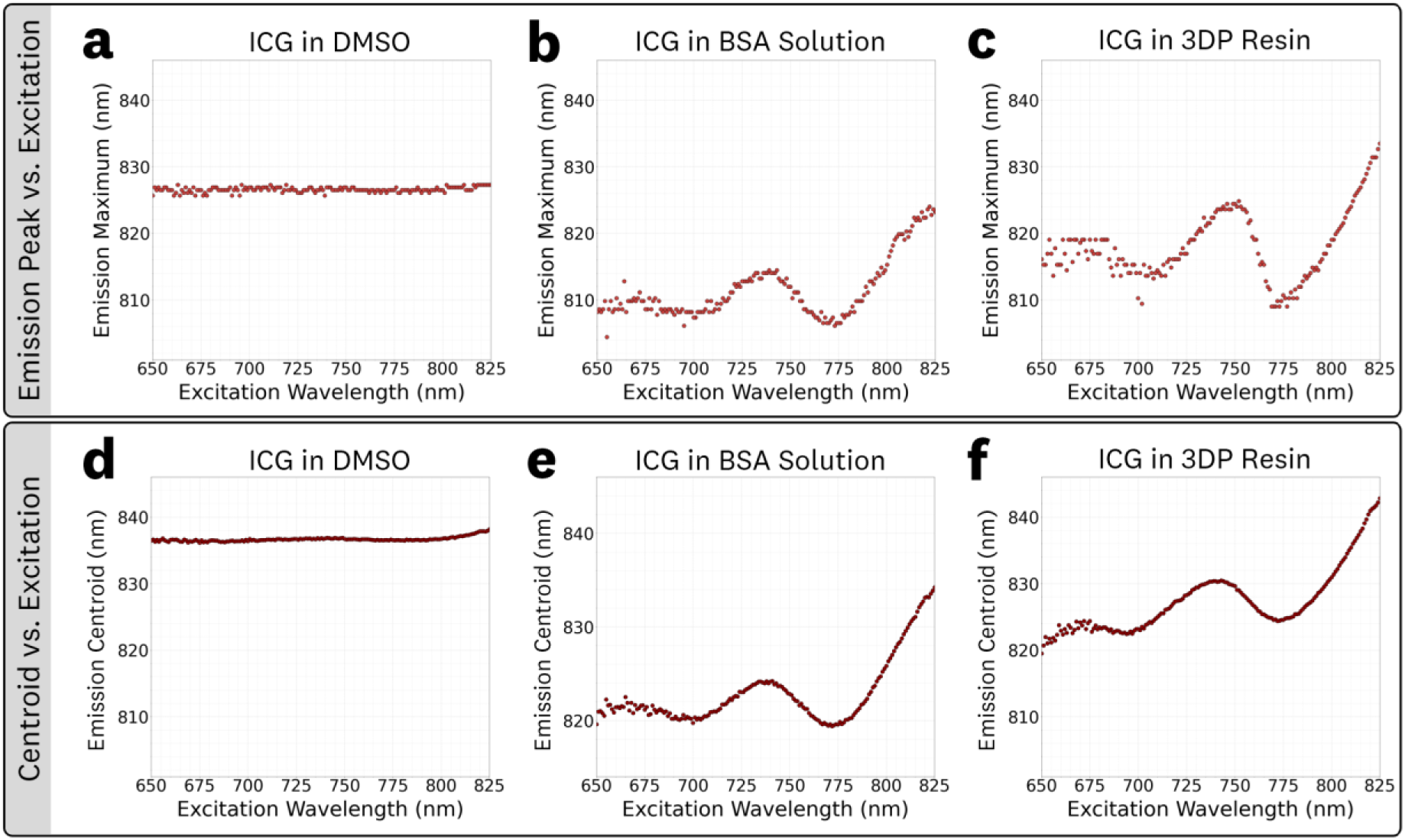
Emission peak **(a-c)** and centroid **(d-f)** vs. excitation wavelength plot of 1 µM ICG in DMSO, BSA solution, and 3DP resin: The peak emission wavelength for **(a)** ICG in DMSO showed no significant variation with excitation wavelength while the **(b)** BSA solution and **(c)** 3DP resin data showed significant, non-linear, variations that include REES. The emission centroid wavelength for **(d)** ICG in DMSO showed no significant variation with excitation wavelength while the **(e)** BSA solution and **(f)** 3DP resin data showed significant, non-linear, variations that include REES.

## 4 Discussion

Here we reported EEM and absorbance measurements of ICG in DMSO, BSA solutions, and 3DP resin, together with extracted excitation and emission spectra and quantitative summaries of emission peaks, centroids, and integrated emission AUCs. ICG in DMSO exhibited excitation-independent emission consistent with Kasha–Vavilov behavior, whereas the BSA solution and 3DP resin showed significant excitation-dependent emission, including REES and non-linear departures from Kasha’s rule. To our knowledge, this is the first report of such excitation-dependent phenomena for ICG and for any FGS fluorophore, highlighting the importance of considering excitation-dependent effects in both comparative spectral analysis and sensing system design. The subsequent subsections provide detailed discussion of each set of results.

### 4.1 ICG in DMSO

The EEM for 1 µM ICG in DMSO (**Fig 3**) provided ‘symmetric’ spectra throughout the varying excitation wavelengths with measured emission peaks of 826.5 nm ± 0.5 nm over the 760 – 810 nm excitation range. This EEM confirms the conventional Kasha-rule fluorophore behavior that assumes invariant spectral photon distribution of fluorescence emission with excitation wavelength.

The EEMs and absorbance at varying ICG concentrations in DMSO (**Fig 4**) show the effects of concentration-dependent fluorescence shifts including fluorescence quenching, CDRS, and disparities between the absorbance and excitation spectra due to IFEs. Increases in fluorophore concentration lead to a proportional increase in fluorescence emission intensity within a linear range, followed by a decrease at higher concentrations due to IFEs or fluorophore aggregation; this concentration-quenching is well understood and reported both experimentally and through Monte Carlo photon transport simulations.^65–67^ The measured linear range for ICG in DMSO was 0.3 – 3 µM with the 10 µM sample showing significant quenching, emission shifts (CDRS) and distortion of the excitation spectra. The 3 µM sample exhibited minor quenching, CDRS, and excitation spectra broadening. Within the linear range, the measured EEMs (**Fig S1**) demonstrated no significant differences, displaying an anticipated increase in emission intensity proportional to concentration across the entire wavelength matrix. The spectral distribution of the absorbance spectra was consistent along the full concentration range, which indicates that the observed fluorescence shifts are due to IFEs and not fluorophore aggregation at these high concentrations. The photon distribution of the absorbance and excitation spectra were equivalent within the identified linear range. It is worth noting that this measured linearity range is specific to the geometry of the spectral measurement, which utilized an orthogonal excitation-emission collection with a 10 mm pathlength. These results highlight the need to understand the linearity range of a fluorophore to adequately choose concentrations for EEM and fluorescence spectra characterization.

### 4.2 ICG in BSA solution

In contrast to the ICG in DMSO sample, the 1 µM ICG in BSA solution (**Fig 5**) showed a ‘rotation’ of the central spectral feature indicating excitation-dependent emission, REES, and departures from Kasha’s rule. This nonlinear effect is most likely attributed to albumin binding^21,68^ causing changes in solvent relaxation times, where the excitation red shift is observed because relaxation is not complete before fluorescence emission occurs.^12^ Quantitatively, average emission red-shift slopes of 0.35 nm·nm⁻¹ (peak) and 0.26 nm·nm⁻¹ (centroid) were observed over the 770–810 nm excitation range (**Table 2**), with only minor changes in spectral photon distributions. These results emphasize the importance of EEM fluorophore measurements to characterize excitation-dependent effects associated with biologically-relevant microenvironments.

EEM measurements for ICG in varying BSA concentration solutions (**Fig 6, Figure S2**) showed the same excitation-dependent behavior, with no significant spectral differences within the biologically-relevant albumin concentrations in the 5–100 mg/mL range.^59,60^ Consideration of excitation-dependent emission shifts for ICG in BSA solutions could help optimize fluorescence sensing designs including FGS imagers given the relevancy to perfusion and *in vivo* applications. The measured ICG EEM in the BSA solution (**Fig 5**) provides spectral information for albumin concentrations found in whole blood,^60,69^ thus serving as a good estimator of *in vivo* ICG spectra for perfusion applications. Furthermore, the invariability of the EEM within the wide albumin concentration range indicates that it might be a suitable estimator of interstitial ICG spectra.^69^

It is worth noting that while higher BSA concentrations introduced greater excitation scattering peaks in the raw data, these effects were successfully addressed by the robust smoothing approach (see Section 2.6), yielding equivalent EEMs and emission spectra across the full concentration range. This demonstrates that the technique can handle varying levels of scattering without negatively affecting the processed results or introducing artifacts within the tested signal-to-noise ratios and smoothing span.

### 4.3 ICG in 3DP resin

Similarly to the ICG in BSA solution EEM, the ICG in 3DP resin EEM (**Fig 7**) showed a ‘rotation’ of the central spectral feature indicating excitation-dependent emission, REES, and departures from Kasha’s rule. Quantitatively, the average emission red-shift slopes were 0.39 nm·nm⁻¹ (peak) and 0.26 nm·nm⁻¹ (centroid) over 770–810 nm, with noticeable changes to spectral photon distribution (broadened spectra) evident from differences between peak-intensity and AUC trends (**Table 3**). These nonlinear effects are most likely attributed to the embedding of the fluorophore within a solid polymer matrix, causing changes in relaxation times, where the excitation red shift is observed because relaxation is not complete before fluorescence emission occurs.^12^ Given the intended use of these 3DP fluorescent materials as ‘ground-truth’ measurements, in the form of reference targets and phantoms,^7,22,23,57^ the presence of excitation-dependent emission underscores the need to consider the excitation wavelength for accurate, application-specific, and system-specific characterization and comparisons.

### 4.4 ICG Spectral comparison

The spectral comparison of 1 µM ICG in DMSO, BSA solution, and 3DP resin (**Fig 8**), showed shifts in spectral emission wavelengths and distributions, including excitation-dependent effects in the BSA and 3DP samples. Given the excitation-dependent spectral behavior, and because fluorescence sensing relies on emitted light, emission spectra at defined excitation wavelengths provide the most appropriate basis for comparison. In this context, the BSA solution and 3DP resin samples exhibited substantial spectral overlap at excitation wavelengths commonly used in FGS devices (760–805 nm, **Fig 8d,f,h**); this suggests that the ICG in 3DP resin can serve as an adequate surrogate reference material for albumin-bound ICG. However, meaningful comparison requires accounting not only for excitation conditions but also for the specific emission collection parameters of the sensing system.

Considering that the fluorophore spectral emission can vary with both microenvironment and excitation wavelength, the most accurate method for comparing fluorescence emission intensity involves integrating the excitation-specific emission spectra within the sensing system’s detection band (such as long-pass emission collection, band-pass emission collection, etc.). The acquired EEM dataset facilitates application and system-specific comparisons, incorporating considerations of both excitation and emission collection parameters. It is important to note that the provided EEMs (Supplementary Data Files) are directly comparable in their reported intensities since identical acquisition parameters were used. Although not explored in our analysis, it is worth noting that ‘weighted emission spectra’ can be generated from the provided EEMs for broadband excitation sources.

### 4.5 Excitation-Dependency and Red-edge Shift Comparisons

The peak and centroid versus excitation plots (**Fig. 9**) showed three distinct behaviors. In DMSO, emission is effectively excitation-independent, following Kasha’s rule, and serves as the baseline for traditional fluorophore behavior assumptions. In BSA solution and 3DP resin, pronounced red-edge excitation shifts emerge for excitations ≥ ∼770 nm, indicating that varied excitation alongside partial vibronic relaxation produce excitation-dependent emission. Additionally, a localized maxima near 740–750 nm precedes the REES trend in both media, pointing to extra microstate selectivity not captured by a simple monotonic shift and representing a more complex departure from Kasha’s rule. The detailed mechanistic explanation for these complex excitation-dependent effects lies beyond the scope of the present work but represents an important direction for further study.

### 4.6 Future work and Limitations

The presented EEM and absorbance measurements provided comprehensive insights into ICG excitation-dependent spectral shifts in varying microenvironments but were limited to room-temperature conditions. It is worth noting that the captured EEMs contain some “banding” along the emission axis (see **Fig. 3a, 5a, 7a**) for excitation wavelengths in the ∼820–860 nm range. These artifacts result from limitations in the spectrofluorometer’s correction of xenon lamp spectral peaks during intensity normalization, as confirmed through communication with the manufacturer. Because the robust smoothing applied to the data was performed independently on each emission spectrum rather than across the excitation axis (the y-axis in the top-down EEM plots), these banding artifacts were not removed. However, the reported peak emissions, centroids, and integrated AUCs, as well as the extracted excitation and emission spectra, were minimally affected, since the relevant excitation range for ICG fluorescence sensing applications lies within the 650– 820 nm range.

Further insights could be gained by including whole blood in the suite of tested solvents. Although the high absorption and scattering in whole blood may pose measurement challenges, understanding excitation-dependence in this context would improve the *in vivo* spectral estimations. Moreover, conducting *in vivo* fluorescence spectral measurements, which would require specialized equipment, could further characterize the effects of albumin binding on *in-vivo* ICG EEM shifts. Characterizing how temperature influences EEMs and excitation-dependent behavior would provide valuable insights. Additionally, IFE corrections^70^ and spectral unmixing of EEMs could further help advance fluorophore characterization, including understanding of non-linear and aggregation effects. Future studies that measure absolute quantum yields at varying excitation wavelengths in BSA solutions and 3DP resin could provide further insight into the extent of deviation from Kasha–Vavilov behavior.

As FGS-targeted fluorophores continue to develop, measuring EEM spectra for bound fluorophores that mimic the *in vivo* environment will be crucial for understanding their behavior and the extent of excitation-dependence and deviations from Kasha’s rule. These assessments are further complicated by the diverse chemical environments of targeted binding sites. If solvent-related excitation effects are found to be minimal, this could indicate ‘robust’ spectral properties, with little to no shift observed despite changes in microenvironment. Conversely, pronounced excitation-dependent spectral changes would be important to identify prior to clinical application, as they could inform imaging system design or motivate modification of the fluorophore structure.

## 5 Conclusion

This study presents comprehensive EEM characterization of ICG in DMSO, BSA solutions, and 3DP resin microenvironments, revealing significant excitation-dependent emission, including REES and non-linear departures from Kasha’s rule, in the BSA solution and 3DP resin; in contrast, DMSO exhibited traditional excitation-independent (Kasha–Vavilov) behavior. These results provide, to our knowledge, the first documentation of REES and broader deviation from Kasha’s rule for ICG and any FGS fluorophore. The findings highlight the need to account for excitation-dependent emission when comparing spectra across systems and microenvironments. The observed excitation-dependent shifts also support the use of excitation-specific, EEM-derived spectra for system design, inter-system comparisons, and phantom development; notably, the substantial emission overlap between BSA solution and 3DP resin spectra at commonly used excitation wavelengths (∼760–805 nm) suggests that ICG in 3DP resin can serve as a stable surrogate reference for albumin-bound ICG when system excitation and emission collection parameters are considered.^7,22,57^ The EEM datasets provided here offer a practical reference for fluorophore characterization and benchmarking that incorporate excitation-dependent effects. Future work extending these measurements to additional physiological conditions (i.e., temperature variation, whole blood, and *in vivo* contexts) and absolute quantum-yield assessments at varying excitations will further clarify excitation-dependent behavior and its implications for fluorophore chemistry, spectral comparisons, and fluorescence-based imaging system design.

### Disclosures

AJR and EPML are co-founders of QUEL Imaging, which is a medical imaging phantom company. SAL and KSS declare no conflicts of interest.

## Supporting information

ICG EEM Data

Loess Python Function

## Acknowledgments

This work was partially funded by the NIBIB R44 grant EB029804-02 and the Dartmouth Innovation Program Fellowship.

The authors also acknowledge the use of the ChatGPT-5 language model (OpenAI) and the Microsoft Word Editor tools for editorial editing, grammar correction, and text refinement.

## Data, Materials, and Code Availability

Data and code are available upon reasonable request.

