## Supplementary Figures for "Indocyanine green excitation-emission matrix characterization: excitation-dependent emission shifts and application-specific spectra"

**Alberto J. Ruiz,[a*] Sophie A. Lyon,[a] Ethan P.M. LaRochelle,[a] Kimberley S. Samkoe,[b]**
[a]QUEL Imaging, White River Junction, VT, USA
[b]Dartmouth Engineering, Hanover, NH, USA

**Table S1:** List of solvents/matrices tested, for which both excitation–emission matrices (EEMs) and absorbance spectra were acquired. The experiment number indicates whether measurements were obtained within the same experimental setup.

| Experiment# | Solvent/Matrix | ICG Concentration (µM) |
|---|---|---|
| 1 | DMSO | 1.00 |
| 1 | DMSO | 0 |
| 1 | BSA – 44 mg/mL | 1.00 |
| 1 | BSA – 44 mg/mL | 0 |
| 1 | 3DP Resin | 1.00 |
| 1 | 3DP Resin | 0 |
| 2 | DMSO | 10.00 |
| 2 | DMSO | 3.00 |
| 2 | DMSO | 1.00 |
| 2 | DMSO | 0.30 |
| 2 | DMSO | 0.10 |
| 2 | DMSO | 0.03 |
| 2 | DMSO | 0 |
| 3 | BSA – 100 mg/mL | 1.00 |
| 3 | BSA – 100 mg/mL | 0 |
| 3 | BSA – 50 mg/mL | 1.00 |
| 3 | BSA – 50 mg/mL | 0 |
| 3 | BSA – 25 mg/mL | 1.00 |
| 3 | BSA – 25 mg/mL | 0 |
| 3 | BSA – 10 mg/mL | 1.00 |
| 3 | BSA – 10 mg/mL | 0 |
| 3 | BSA – 5 mg/mL | 1000 |
| 3 | BSA – 5 mg/mL | 0 |

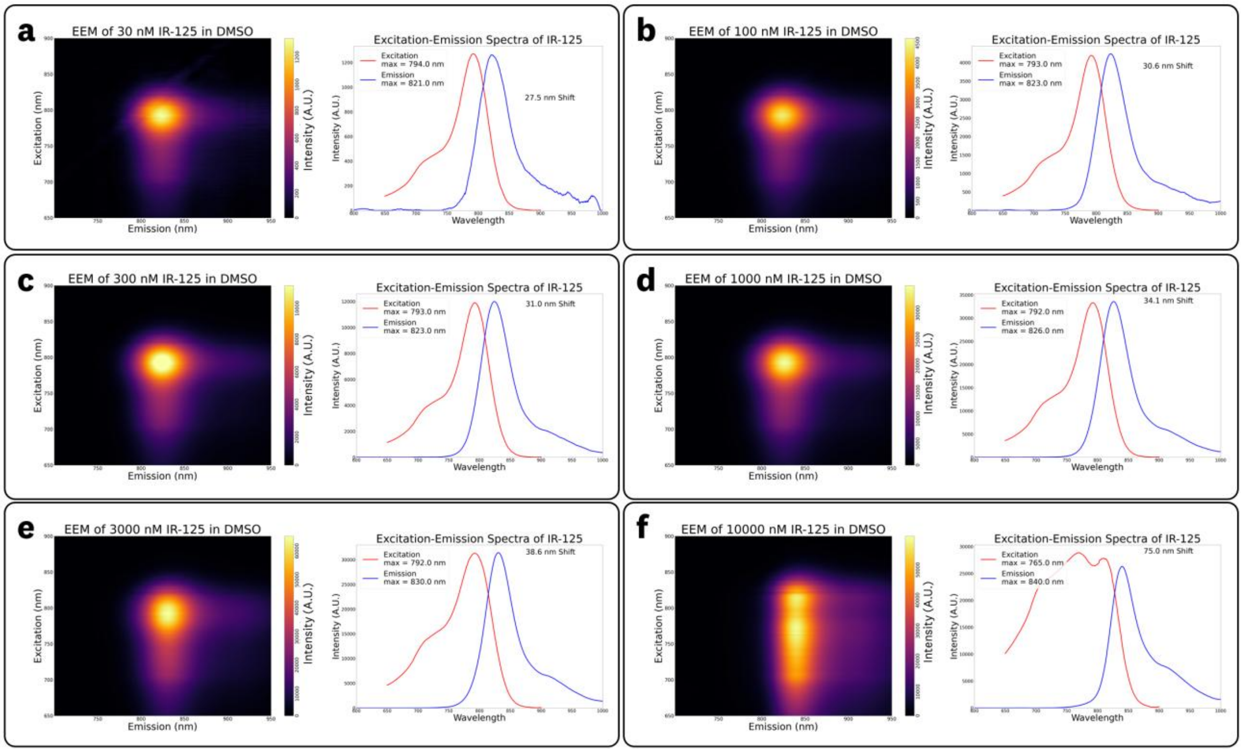


**Figure S1:** EEMs for ICG (IR-125) in DMSO for varying concentrations alongside excitation-emission spectra at the EEM maxima for (a) 30 nM, (b) 100 nM, (c) 300 nM, (d) 1000 nM, (e) 3000 nM, and (f) 10,000 nM.

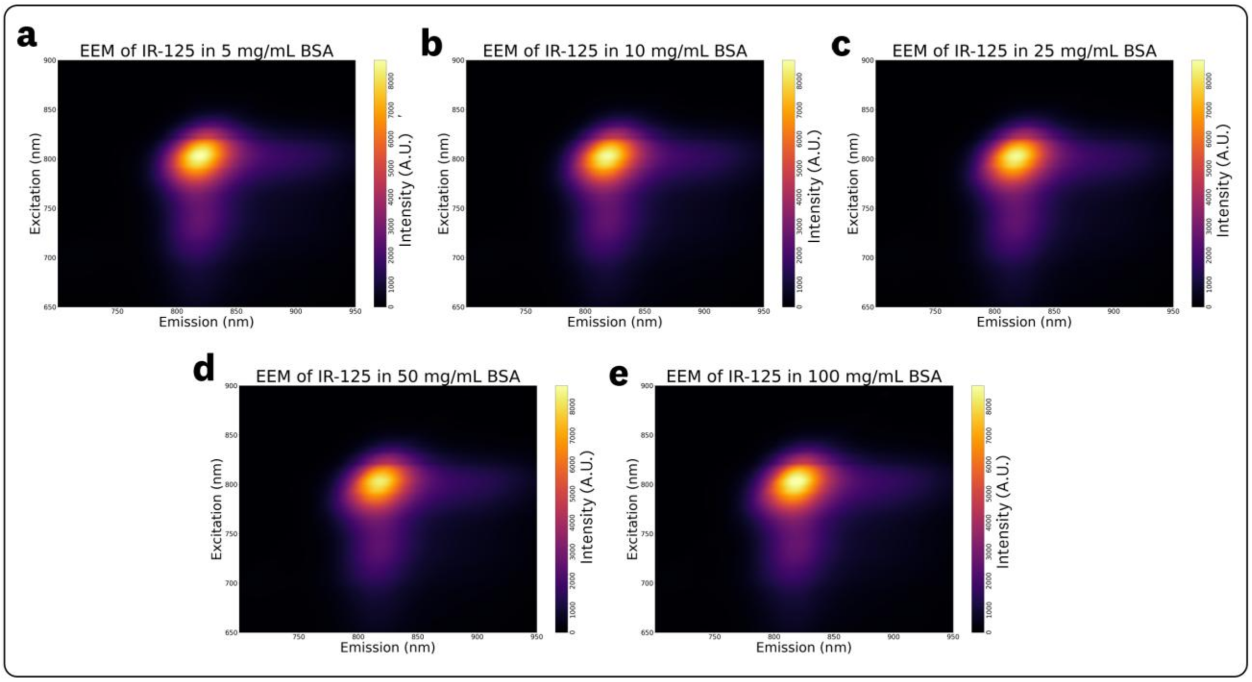


**Figure S2**: EEMS for ICG (IR-125) in varying BSA solution concentrations of (a) 5 mg/mL, (b) 10 mg/mL, (c) 25 mg/mL, (d) 50 mg/mL, and (e) 100 mg/mL.

**Table S2**: Absorbance data for varying ICG concentrations in DMSO. Note that the 0.03 µM data is provided for completeness but should be considered an outlier due to the low signal-to-noise associated with the measurement.

| Concentration (µM) | Max Wavelength (nm) | Max Absorbance (OD) | Absorbance at 785 nm (OD) | Absorbance at 805 nm (OD) | Absorbance at 825 nm (OD) |
|---|---|---|---|---|---|
| 0.03 | 798 | 0.0110 | 0.0103 | 0.0108 | 0.0089 |
| 0.1 | 794 | 0.0179 | 0.0168 | 0.0158 | 0.0068 |
| 0.3 | 794 | 0.0578 | 0.0544 | 0.0519 | 0.0257 |
| 1 | 793 | 0.1853 | 0.1744 | 0.1642 | 0.0754 |
| 3 | 793 | 0.5503 | 0.5188 | 0.4833 | 0.2156 |
| 10 | 793 | 1.9113 | 1.7898 | 1.6529 | 0.7073 |

**Table S3**: Calculated molar extinction coefficients (ε) for varying ICG concentrations in DMSO based on Table S1 results. Note that the 0.03 µM calculation is provided for completeness but should be considered an outlier due to the low signal-to-noise associated with the measurement.

| Concentration (µM) | Max Wavelength (nm) | Calculated ε at max abs ($\mu M^{-1}$ $cm^{-1}$) | Calculated ε at 785 nm ($\mu M^{-1}$ $cm^{-1}$) | Calculated ε at 805 nm ($\mu M^{-1}$ $cm^{-1}$) | Calculated ε at 825 nm ($\mu M^{-1}$ $cm^{-1}$) |
|---|---|---|---|---|---|
| 0.03 | 798 | 0.3677 | 0.3437 | 0.3601 | 0.2954 |
| 0.1 | 794 | 0.1790 | 0.1683 | 0.1581 | 0.0678 |
| 0.3 | 794 | 0.1925 | 0.1812 | 0.1730 | 0.0856 |
| 1 | 793 | 0.1853 | 0.1744 | 0.1642 | 0.0754 |
| 3 | 793 | 0.1834 | 0.1729 | 0.1611 | 0.0719 |
| 10 | 793 | 0.1911 | 0.1790 | 0.1653 | 0.0707 |
