## Supplementary material for "Indocyanine green excitation-emission matrix characterization: excitation-dependent emission shifts and application-specific spectra": Loess Python Function: METADATA

Metadata-Version: 2.1
Name: loess
Version: 2.1.2
Summary: LOESS: smoothing via robust locally-weighted regression in one or two dimensions
Home-page: https://purl.org/cappellari/software
Author: Michele Cappellari
Author-
License: Other/Proprietary License
Classifier: Development Status :: 5 - Production/Stable
Classifier: Intended Audience :: Developers
Classifier: Intended Audience :: Science/Research
Classifier: Operating System :: OS Independent
Classifier: Programming Language :: Python :: 3
Description-Content-Type: text/x-rst
Requires-Dist: numpy
Requires-Dist: matplotlib
Requires-Dist: plotbin

The LOESS Package
=================

**Smoothing via robust locally-weighted regression in one or two dimensions**

.. image:: https://img.shields.io/pypi/v/loess.svg
        :target: https://pypi.org/project/loess/
.. image:: https://img.shields.io/badge/arXiv-1208.3523-orange.svg
    :target: https://arxiv.org/abs/1208.3523
.. image:: https://img.shields.io/badge/DOI-10.1093/mnras/stt644-green.svg
        :target: https://doi.org/10.1093/mnras/stt644

LOESS is the Python implementation by `Cappellari et al. (2013)
<https://ui.adsabs.harvard.edu/abs/2013MNRAS.432.1862C>`_ of the
algorithm by `Cleveland (1979) <https://doi.org/10.2307/2286407>`_
for the one-dimensional case and `Cleveland & Devlin (1988)
<https://doi.org/10.2307/2289282>`_ for the two-dimensional case.

.. contents:: :depth: 2

Attribution
-----------

If you use this software for your research, please cite the LOESS package of
`Cappellari et al. (2013b) <https://ui.adsabs.harvard.edu/abs/2013MNRAS.432.1862C>`_,
where the implementation was described. The BibTeX entry for the paper is::

    @ARTICLE{Cappellari2013b,
        author = {{Cappellari}, M. and {McDermid}, R.~M. and {Alatalo}, K. and 
            {Blitz}, L. and {Bois}, M. and {Bournaud}, F. and {Bureau}, M. and 
            {Crocker}, A.~F. and {Davies}, R.~L. and {Davis}, T.~A. and 
            {de Zeeuw}, P.~T. and {Duc}, P.-A. and {Emsellem}, E. and {Khochfar}, S. and 
            {Krajnovi{\'c}}, D. and {Kuntschner}, H. and {Morganti}, R. and 
            {Naab}, T. and {Oosterloo}, T. and {Sarzi}, M. and {Scott}, N. and 
            {Serra}, P. and {Weijmans}, A.-M. and {Young}, L.~M.},
        title = "{The ATLAS$^{3D}$ project - XX. Mass-size and mass-{$\sigma$}
            distributions of early-type galaxies: bulge fraction drives kinematics,
            mass-to-light ratio, molecular gas fraction and stellar initial mass
            function}",
        journal = {MNRAS},
        eprint = {1208.3523},
         year = 2013,
        volume = 432,
        pages = {1862-1893},
          doi = {10.1093/mnras/stt644}
    }

Installation
------------

install with::

    pip install loess

Without writing access to the global ``site-packages`` directory, use::

    pip install --user loess

To upgrade ``loess`` to the latest version use::

    pip install --upgrade loess

Documentation
-------------

Full documentation is contained in the individual files docstrings.

Usage examples are contained in the directory ``loess/examples`` 
which is copied by ``pip`` within the global folder
`site-packages <https://stackoverflow.com/a/46071447>`_.

What follows is the documentation of the two main procedures of the 
``loess`` package, extracted from their Python docstrings.

###########################################################################

loess_1d
========

Purpose
-------

One-dimensional LOESS smoothing via robust locally-weighted regression.

This function is the implementation by `Cappellari et al. (2013)
<https://ui.adsabs.harvard.edu/abs/2013MNRAS.432.1862C>`_ of the
algorithm by `Cleveland (1979) <https://doi.org/10.2307/2286407>`_.

Calling Sequence
----------------

.. code-block:: python

    xout, yout, wout = loess_1d(x, y, xnew=None, degree=1, frac=0.5,
                                npoints=None, rotate=False, sigy=None)

Input Parameters
----------------

x: array_like with shape (n,)
    Vector of ``x`` coordinate.
y: array_like with shape (n,)
    Vector of ``y`` coordinate to be LOESS smoothed.

Optional Keywords
-----------------

xnew: array_like with shape (m,), optional
    Vector of coordinates at which to compute the smoothed ``y`` values.
degree: {1, 2}, optional
    degree of the local 1-dim polynomial approximation (default ``degree=1``).
frac: float, optional
    Fraction of points to consider in the local approximation (default ``frac=0.5``).
    Typical values are between ``frac~0.2-0.8``. Note that the values are
    weighted by a Gaussian function of their distance from the point under 
    consideration. This implies that the effective fraction of points 
    contributing to a given value is much smaller that ``frac``.
npoints: int, optional
    Number of points to consider in the local approximation.
    This is an alternative to using ``frac=npoints/x.size``.
rotate: bool, optional
    Rotate the ``(x, y)`` coordinates to have the maximum variance along the
    ``x`` axis. This is useful to give comparable contribution to the
    errors in the ``x`` and ``y`` variables. It can be used to asses the
    sensitivity of the solution to the assumption that errors are only in ``y``.
sigy: array_like with shape (n,)
    1-sigma errors for the ``y`` values. If this keyword is used
    the biweight fit is done assuming those errors. If this keyword
    is *not* used, the biweight fit determines the errors in ``y``
    from the scatter of the neighbouring points.

Output Parameters
-----------------

xout: array_like with shape (n,)
    Vector of ``x`` coordinates for the ``yout`` values.
    If ``rotate=False`` (default) then ``xout=x``.
    
    When passing as input the ``xnew`` coordinates then ``xout=xnew``
    and both have shape ``(m,)``.
yout: array_like with shape (n,)
    Vector of smoothed ``y`` values at the coordinates ``xout``.
    
    When passing as input the ``xnew`` coordinates this contains the
    smoothed values at the coordinates ``xnew`` and has shape ``(m,)``.
wout: array_like with shape (n,)
    Vector of biweights used in the local regressions. This can be used to
    identify outliers: ``wout=0`` for outliers with deviations ``>4sigma``.
    
    When passing as input the ``xnew`` coordinates, this output is
    meaningless and is arbitrarily set to unity.

###########################################################################

loess_2d
========

Purpose
-------

Two-dimensional LOESS smoothing via robust locally-weighted regression.

This function is the implementation by `Cappellari et al. (2013)
<https://ui.adsabs.harvard.edu/abs/2013MNRAS.432.1862C>`_ of the
algorithm by `Cleveland (1979) <https://doi.org/10.2307/2286407>`_
for the one-dimensional case and `Cleveland & Devlin (1988)
<https://doi.org/10.2307/2289282>`_ for the two-dimensional case.

Calling Sequence
----------------

.. code-block:: python

   zout, wout = loess_2d(x, y, z, xnew=None, ynew=None, degree=1, frac=0.5,
                         npoints=None, rescale=False, sigz=None)

Input Parameters
----------------

x: array_like with shape (n,)
    vector of ``x`` coordinates.
y: array_like with shape (n,)
    vector of ``y`` coordinates.
z: array_like with shape (n,)
    vector of ``z`` coordinates to be LOESS smoothed.

Optional Keywords
-----------------

xnew: array_like with shape (m,), optional
    Vector with the ``x`` coordinates at which to compute the smoothed
    ``z`` values.
ynew: array_like with shape (m,), optional
    Vector with the ``y`` coordinates at which to compute the smoothed
    ``z`` values.
degree: {1, 2}, optional
    degree of the local 2-dim polynomial approximation (default ``degree=1``).
frac: float, optional
    Fraction of points to consider in the local approximation (default ``frac=0.5``).
    Typical values are between ``frac~0.2-0.8``. Note that the values are
    weighted by a Gaussian function of their distance from the point under
    consideration. This implies that the effective fraction of points
    contributing to a given value is much smaller that ``frac``.
npoints: int, optional
    Number of points to consider in the local approximation.
    This is an alternative to using ``frac=npoints/x.size``.
rescale: bool, optional
    Rotate the ``(x, y)`` coordinates to make the ``x`` axis the axis of
    maximum variance. Subsequently scale the coordinates to have equal
    variance along both axes. Then perform the local regressions.
    This is recommended when the distribution of points is elongated or
    when the units are very different for the two axes.
sigz: array_like with shape (n,)
    1-sigma errors for the ``z`` values. If this keyword is used
    the biweight fit is done assuming these errors. If this keyword
    is *not* used, the biweight fit determines the errors in ``z``
    from the scatter of the neighbouring points.

Output Parameters
-----------------

zout: array_like with shape (n,)
    Vector of smoothed ``z`` values at the coordinates ``(x, y)``, or at
    ``(xnew, ynew)`` if the latter are given as input. In the latter case
    ``zout`` has shape ``(m,)``.
wout: array_like with shape (n,)
    Vector of biweights used in the local regressions. This can be used to
    identify outliers: ``wout=0`` for outliers with deviations ``>4sigma``.

    When passing as input the ``(xnew, ynew)`` coordinates, this output is
    meaningless and is arbitrarily set to unity.

###########################################################################


License
=======

Other/Proprietary License

Copyright (c) 2010-2022 Michele Cappellari

This software is provided as is without any warranty whatsoever.
Permission to use, for non-commercial purposes is granted.
Permission to modify for personal or internal use is granted,
provided this copyright and disclaimer are included in all 
copies of the software. All other rights are reserved.
In particular, redistribution of the code is not allowed.

Changelog
=========

V2.1.2: MC, Oxford, 31 January 2022
    - Fixed incorrect results with integer input coordinates in both
      ``loess_1d`` and ``loess_2d``. Thanks to Peter Weilbacher (aip.de) 
      for the report and fix.

V2.1.0: MC, Oxford, 20 July 2021 
    - Support output coordinates different from the input ones.
    - Updated ``loess_1d_example`` and ``loess_2d_example``.
       
V2.0.6: MC, Oxford, 21 May 2018
    - Dropped support for Python 2.7. 

V2.0.5: MC, Oxford, 18 January 2018
    - Fixed FutureWarning in Numpy 1.14. 

V2.0.4: MC, Oxford, 18 April 2016
    - Fixed deprecation warning in Numpy 1.11. 

V2.0.3: MC, Oxford, 8 December 2014
    - Updated documentation. Minor polishing. 

V2.0.2: MC, Oxford, 3 November 2014
    - Returns weights also when frac=0 for consistency.

V2.0.1: MC, Oxford, 10 July 2014
    - Removed SciPy dependency. 
   
V2.0.0: MC, Oxford, 26 February 2014
    - Translated from IDL into Python. 

V1.3.4: MC, Paranal, 7 November 2013
    - Include SIGZ and WOUT keywords. Updated documentation.
   
V1.3.3: MC, Oxford, 31 October 2013
    - Use CAP_POLYFIT_2D. 
    - Removed /QUARTIC keyword and replaced by DEGREE keyword like CAP_LOESS_1D.

V1.3.2: MC, Oxford, 12 October 2013
    - Test whether input (X,Y,Z) have the same size.
    - Included NPOINTS keyword. 

V1.1.4: MC, Oxford, 16 May 2013
    - Updated documentation. 

V1.1.3: MC, Oxford, 2 December 2011
    - Check when outliers don't change to stop iteration.
   
V1.1.2: MC, Oxford, 25 July 2011
    - Return values unchanged if FRAC=0. 

V1.1.1: MC, Oxford, 07 March 2011 
    - Fix: use ABS() for proper computation of "r".
   
V1.1.0: MC, Vicenza, 30 December 2010 
    - Rescale after rotating to axis of maximum variance.
   
V1.0.0: Michele Cappellari, Oxford, 15 December 2010
    - Written and tested.
   
