## Supplementary material for "Indocyanine green excitation-emission matrix characterization: excitation-dependent emission shifts and application-specific spectra": Loess Python Function: loess_1d_example.py

```
#!/usr/bin/env python
##############################################################################
#
# Usage example for the procedure loess_1d
#
# MODIFICATION HISTORY:
#   V1.0.0: Written by Michele Cappellari, Oxford 25 February 2015
#   V1.0.1: Fixed deprecation warning in Numpy 1.11. MC, Oxford, 18 April 2016
#   V1.0.2: Changed imports for loess as a package. MC, Oxford, 17 April 2018
#   V1.1.0: Show how to sample the output values at a different set of coordinates.
#       MC, Oxford, 20 July 2021
#
##############################################################################

import numpy as np
import matplotlib.pyplot as plt

from loess.loess_1d import loess_1d

def loess_1d_example():
    """ Usage example for loess_1d """

    n = 50
    nbad = int(n*0.1) # 10% outliers
    np.random.seed(5)

    # Compute the true model
    xtrue = np.random.uniform(-np.pi, np.pi, n - nbad)
    xtrue.sort()   # Sort only for plotting smooth lines
    ytrue = np.sin(xtrue) + xtrue/2

    # Add noise to the data
    sigy = 0.5
    yran = np.random.normal(ytrue, sigy)

    # Add outliers to the data
    xbad = np.random.uniform(-np.pi, np.pi, nbad)
    ybad = np.random.uniform(-5, 5, nbad)
    xfit = np.append(xtrue, xbad)
    yfit = np.append(yran, ybad)

    # Sort only for plotting smooth lines
    j = np.argsort(xfit)
    xfit, yfit = xfit[j], yfit[j]

    # LOESS smoothing on the same coordinates as input
    xout1, yout1, weights1 = loess_1d(xfit, yfit, frac=0.8, degree=2)

    # LOESS smoothing on a new output grid (xnew)
    xnew = np.linspace(np.min(xfit), np.max(xfit), 30)
    xout2, yout2, weights2 = loess_1d(xfit, yfit, xnew, frac=0.8, degree=2)

    plt.clf()
    plt.subplot(211)
    plt.plot(xfit, yfit, 'ro', label='Noisy')
    plt.plot(xtrue, ytrue, color='limegreen', label='True')
    plt.plot(xout1, yout1, '+-b', label='LOESS')
    w = weights1 < 0.34  # identify outliers
    plt.plot(xfit[w], yfit[w], 'xk', ms=10, label='Outliers')
    plt.title("LOESS smoothing at the input coordinates")
    plt.legend()

    plt.subplot(212)
    plt.plot(xfit, yfit, 'ro', label='Noisy')
    plt.plot(xtrue, ytrue, color='limegreen', label='True')
    plt.plot(xout2, yout2, '+-b', label='LOESS')
    plt.title("LOESS smoothing on a regular output grid")
    plt.legend()

    plt.tight_layout()
    plt.pause(1)

#-----------------------------------------------------------------------------

if __name__ == '__main__':

    loess_1d_example()
```
