## Supplementary material for "Indocyanine green excitation-emission matrix characterization: excitation-dependent emission shifts and application-specific spectra": Loess Python Function: loess_2d_example.py

```
#!/usr/bin/env python
##############################################################################
#
# Usage example for the procedure loess_2d
#
# MODIFICATION HISTORY:
#   V1.0.0: Written by Michele Cappellari, Oxford 26 February 2014
#   V1.0.1: Changed imports for loess and plotbin as packages. 
#       Removed overlap of axes labels. MC, Oxford, 17 April 2018
#   V1.0.2: Fixed clock DeprecationWarning in Python 3.7.
#       MC, Oxford, 27 September 2018
#   V1.1.0: Show how to sample the output values at a different set of coordinates.
#       MC, Oxford, 20 July 2021
#
##############################################################################

import numpy as np
import matplotlib.pyplot as plt

from loess.loess_2d import loess_2d
from plotbin.plot_velfield import plot_velfield

def loess_2d_example():
    """ Usage example for loess_2d """

    n = 200
    np.random.seed(1)

    # Compute the true model
    x, y = np.random.uniform(-1, 1, size=[n, 2]).T
    z = x**2 - y**2

    # Add noise to the data
    sigz = 0.2
    zran = np.random.normal(z, sigz)

    # LOESS smoothing on the same coordinates as input
    zout1, wout = loess_2d(x, y, zran)

    # LOESS smoothing on a new output grid (xnew, ynew)
    xx = np.linspace(np.min(x), np.max(x), 15)
    yy = np.linspace(np.min(y), np.max(y), 15)
    xnew, ynew = map(np.ravel, np.meshgrid(xx, yy))
    zout2, wout = loess_2d(x, y, zran, xnew, ynew)

    plt.clf()
    plt.subplot(221)
    plot_velfield(x, y, z)
    plt.title("True Function")

    plt.subplot(222)
    plot_velfield(x, y, zran)
    plt.title("With Noise Added")

    plt.subplot(223)
    plot_velfield(x, y, zout1)
    plt.title("LOESS Recovery Input Grid")

    plt.subplot(224)
    plot_velfield(xnew, ynew, zout2, markersize=1)
    plt.title("LOESS Recovery New Grid")

    plt.tight_layout()
    plt.pause(1)

#------------------------------------------------------------------------

if __name__ == '__main__':

    loess_2d_example()
```
