## Supplementary material for "Indocyanine green excitation-emission matrix characterization: excitation-dependent emission shifts and application-specific spectra": Loess Python Function: loess_1d.py

```
"""

    Copyright (C) 2010-2022, Michele Cappellari
    E-mail: michele.cappellari_at_physics.ox.ac.uk

    Updated versions of the software are available from my web page
    http://purl.org/cappellari/software

    If you have found this software useful for your research,
    I would appreciate an acknowledgement to the use of the
    "LOESS_1D routine of Cappellari et al. (2013b), which
    implements the univariate LOESS algorithm of Cleveland (1979)"

    https://ui.adsabs.harvard.edu/abs/2013MNRAS.432.1862C

    This software is provided as is without any warranty whatsoever.
    Permission to use, for non-commercial purposes is granted.
    Permission to modify for personal or internal use is granted,
    provided this copyright and disclaimer are included unchanged
    at the beginning of the file. All other rights are reserved.
    In particular, redistribution of the code is not allowed.

Changelog
---------

   V1.0.0: Michele Cappellari Oxford, 15 December 2010
   V1.1.0: Rescale after rotating to axis of maximum variance.
       MC, Vicenza, 30 December 2010
   V1.1.1: Fix: use ABS() for proper computation of "r".
       MC, Oxford, 07 March 2011
   V1.1.2: Return values unchanged if FRAC=0. MC, Oxford, 25 July 2011
   V1.1.3: Check when outliers don't change to stop iteration.
       MC, Oxford, 2 December 2011
   V1.1.4: Updated documentation. MC, Oxford, 16 May 2013
   V1.3.2: Test whether input (X,Y,Z) have the same size.
       Included NPOINTS keyword. MC, Oxford, 12 October 2013
   V1.3.3: Use CAP_POLYFIT_2D. Removed /QUARTIC keyword and replaced
       by DEGREE keyword like CAP_LOESS_1D. MC, Oxford, 31 October 2013
   V1.3.4: Include SIGZ and WOUT keywords. Updated documentation.
       MC, Paranal, 7 November 2013
   V2.0.0: Translated from IDL into Python. MC, Oxford, 26 February 2014
   V2.0.1: Removed SciPy dependency. MC, Oxford, 10 July 2014
   V2.0.2: Returns weights also when frac=0 for consistency.
       MC, Oxford, 3 November 2014
   V2.0.3: Updated documentation. Minor polishing. MC, Oxford, 8 December 2014
   V2.0.4: Converted from 2D to 1D. MC, Oxford, 23 February 2015
   V2.0.5: Updated documentation. MC, Oxford, 26 March 2015
   V2.0.6: Fixed deprecation warning in Numpy 1.11. MC, Oxford, 18 April 2016
   V2.0.7: Allow polyfit_1d() to be called without errors/weights.
       MC, Oxford, 10 February 2017
   V2.0.8: Fixed FutureWarning in Numpy 1.14. MC, Oxford, 18 January 2018
   V2.0.9: Dropped support for Python 2.7. MC, Oxford, 21 May 2018
   V2.1.0: Allow one to specify output coordinates different from the input ones.
      MC, Oxford, 20 July 2021
   Vx.x.x: additional changes are documented in the global package CHANGELOG.

"""
import numpy as np


################################################################################


class polyfit1d:

    def __init__(self, x, y, degree, weights):
        """
        Fit a univariate polynomial of given DEGREE to a set of points
        (X, Y), assuming errors in the Y variable only and weights=1/sigy^2.

        For example with DEGREE=1 this function fits a straight line

           y = a + b*x

        while with DEGREE=2 the function fits a parabola

           y = a + b*x + c*x^2

        """
        sqw = np.sqrt(weights)
        a = x[:, None]**np.arange(degree + 1)
        self.degree = degree
        self.coeff = np.linalg.lstsq(a*sqw[:, None], y*sqw, rcond=None)[0]
        self.yfit = a @ self.coeff


    def eval(self, x):
        """Evaluate at the coordinate x the polynomial previously fitted"""

        a = x**np.arange(self.degree + 1)
        yout = a @ self.coeff

        return yout


################################################################################


def biweight_sigma(y, zero=False):
    """
    Biweight estimate of the scale (standard deviation).
    Implements the approach described in
    "Understanding Robust and Exploratory Data Analysis"
    Hoaglin, Mosteller, Tukey ed., 1983, Chapter 12B, pg. 417

    """
    y = np.ravel(y)
    if zero:
        d = y
    else:
        d = y - np.median(y)

    mad = np.median(np.abs(d))
    u2 = (d / (9.*mad))**2  # c = 9
    good = u2 < 1.
    u1 = 1. - u2[good]
    num = y.size * ((d[good]*u1**2)**2).sum()
    den = (u1*(1. - 5.*u2[good])).sum()
    sigma = np.sqrt(num/(den*(den - 1.)))  # see note in above reference

    return sigma


################################################################################


def rotate_points(x, y, ang):
    """
    Rotates points counter-clockwise by an angle ANG in degrees.
    Michele cappellari, Paranal, 10 November 2013

    """
    theta = np.radians(ang)
    xNew = x*np.cos(theta) - y*np.sin(theta)
    yNew = x*np.sin(theta) + y*np.cos(theta)

    return xNew, yNew


################################################################################


def loess_1d(x, y, xnew=None, degree=1, frac=0.5, npoints=None, rotate=False, sigy=None):
    """
    loess_1d
    ========

    Purpose
    -------

    One-dimensional LOESS smoothing via robust locally-weighted regression.

    This function is the implementation by `Cappellari et al. (2013)
    <https://ui.adsabs.harvard.edu/abs/2013MNRAS.432.1862C>`_ of the
    algorithm by `Cleveland (1979) <https://doi.org/10.2307/2286407>`_.

    Calling Sequence
    ----------------

    .. code-block:: python

        xout, yout, wout = loess_1d(x, y, xnew=None, degree=1, frac=0.5,
                                    npoints=None, rotate=False, sigy=None)

    Input Parameters
    ----------------

    x: array_like with shape (n,)
        Vector of ``x`` coordinate.
    y: array_like with shape (n,)
        Vector of ``y`` coordinate to be LOESS smoothed.

    Optional Keywords
    -----------------

    xnew: array_like with shape (m,), optional
        Vector of coordinates at which to compute the smoothed ``y`` values.
    degree: {1, 2}, optional
        degree of the local 1-dim polynomial approximation (default ``degree=1``).
    frac: float, optional
        Fraction of points to consider in the local approximation (default ``frac=0.5``).
        Typical values are between ``frac~0.2-0.8``. Note that the values are
        weighted by a Gaussian function of their distance from the point under 
        consideration. This implies that the effective fraction of points 
        contributing to a given value is much smaller that ``frac``.
    npoints: int, optional
        Number of points to consider in the local approximation.
        This is an alternative to using ``frac=npoints/x.size``.
    rotate: bool, optional
        Rotate the ``(x, y)`` coordinates to have the maximum variance along the
        ``x`` axis. This is useful to give comparable contribution to the
        errors in the ``x`` and ``y`` variables. It can be used to asses the
        sensitivity of the solution to the assumption that errors are only in ``y``.
    sigy: array_like with shape (n,)
        1-sigma errors for the ``y`` values. If this keyword is used
        the biweight fit is done assuming those errors. If this keyword
        is *not* used, the biweight fit determines the errors in ``y``
        from the scatter of the neighbouring points.

    Output Parameters
    -----------------

    xout: array_like with shape (n,)
        Vector of ``x`` coordinates for the ``yout`` values.
        If ``rotate=False`` (default) then ``xout=x``.
        
        When passing as input the ``xnew`` coordinates then ``xout=xnew``
        and both have shape ``(m,)``.
    yout: array_like with shape (n,)
        Vector of smoothed ``y`` values at the coordinates ``xout``.
        
        When passing as input the ``xnew`` coordinates this contains the
        smoothed values at the coordinates ``xnew`` and has shape ``(m,)``.
    wout: array_like with shape (n,)
        Vector of biweights used in the local regressions. This can be used to
        identify outliers: ``wout=0`` for outliers with deviations ``>4sigma``.
        
        When passing as input the ``xnew`` coordinates, this output is
        meaningless and is arbitrarily set to unity.

    ###########################################################################
    """

    if frac == 0:
        return y, np.ones_like(y)

    assert x.size == y.size, 'Input vectors (X, Y) must have the same size'

    if npoints is None:
        npoints = int(np.ceil(frac*x.size))

    if rotate:

        assert xnew is None, "`rotate` not supported with `xnew`"

        # Robust calculation of the axis of maximum variance
        #
        nsteps = 180
        angles = np.arange(nsteps)
        sig = np.zeros(nsteps)
        for j, ang in enumerate(angles):
            x2, y2 = rotate_points(x, y, ang)
            sig[j] = biweight_sigma(x2)
        k = np.argmax(sig)  # Find index of max value
        x, y = rotate_points(x, y, angles[k])

    if xnew is None:
        xnew = x

    ynew = np.empty_like(xnew, dtype=float)
    wout = np.empty_like(ynew)

    for j, xj in enumerate(xnew):

        dist = np.abs(x - xj)
        w = np.argsort(dist)[:npoints]
        dist_weights = (1 - (dist[w]/dist[w[-1]])**3)**3  # tricube function distance weights
        yfit = polyfit1d(x[w], y[w], degree, dist_weights).yfit

        # Robust fit from Sec.2 of Cleveland (1979)
        # Use errors if those are known.
        #
        bad = None
        for p in range(10):  # do at most 10 iterations
        
            if sigy is None:                # Errors are unknown
                aerr = np.abs(yfit - y[w])  # Note ABS()
                mad = np.median(aerr)       # Characteristic scale
                uu = (aerr/(6*mad))**2      # For a Gaussian: sigma=1.4826*MAD
            else:                           # Errors are assumed known
                uu = ((yfit - y[w])/(4*sigy[w]))**2  # 4*sig ~ 6*mad

            uu = uu.clip(0, 1)
            biweights = (1 - uu)**2
            tot_weights = dist_weights*biweights
            poly = polyfit1d(x[w], y[w], degree, tot_weights)
            yfit = poly.yfit
            badOld = bad
            bad = biweights < 0.34    # 99% confidence outliers
            if np.array_equal(badOld, bad):
                break

        if np.array_equal(x, xnew):
            ynew[j] = yfit[0]
            wout[j] = biweights[0]
        else:
            ynew[j] = poly.eval(xj)
            wout[j] = 1

    if rotate:
        xnew, ynew = rotate_points(xnew, ynew, -angles[k])
        j = np.argsort(xnew)
        xnew, ynew = xnew[j], ynew[j]

    return xnew, ynew, wout

################################################################################
```
