## Supplementary material for "Indocyanine green excitation-emission matrix characterization: excitation-dependent emission shifts and application-specific spectra": Loess Python Function: loess_2d.py

```
"""
    Copyright (C) 2010-2022, Michele Cappellari
    E-mail: michele.cappellari_at_physics.ox.ac.uk

    Updated versions of the software are available from my web page
    http://purl.org/cappellari/software

    If you have found this software useful for your research,
    I would appreciate an acknowledgement to the use of the
    "LOESS_2D routine of Cappellari et al. (2013b), which implements
    the multivariate LOESS algorithm of Cleveland & Devlin (1988)"

    https://ui.adsabs.harvard.edu/abs/2013MNRAS.432.1862C

    This software is provided as is without any warranty whatsoever.
    Permission to use, for non-commercial purposes is granted.
    Permission to modify for personal or internal use is granted,
    provided this copyright and disclaimer are included unchanged
    at the beginning of the file. All other rights are reserved.
    In particular, redistribution of the code is not allowed.

Changelog
---------

   V1.0.0: Michele Cappellari Oxford, 15 December 2010
   V1.1.0: Rescale after rotating to axis of maximum variance.
       MC, Vicenza, 30 December 2010
   V1.1.1: Fix: use ABS() for proper computation of "r".
       MC, Oxford, 07 March 2011
   V1.1.2: Return values unchanged if FRAC=0. MC, Oxford, 25 July 2011
   V1.1.3: Check when outliers don't change to stop iteration.
       MC, Oxford, 2 December 2011
   V1.1.4: Updated documentation. MC, Oxford, 16 May 2013
   V1.3.2: Test whether input (X,Y,Z) have the same size.
       Included NPOINTS keyword. MC, Oxford, 12 October 2013
   V1.3.3: Use CAP_POLYFIT_2D. Removed /QUARTIC keyword and replaced
       by DEGREE keyword like CAP_LOESS_1D. MC, Oxford, 31 October 2013
   V1.3.4: Include SIGZ and WOUT keywords. Updated documentation.
       MC, Paranal, 7 November 2013
   V2.0.0: Translated from IDL into Python. MC, Oxford, 26 February 2014
   V2.0.1: Removed SciPy dependency. MC, Oxford, 10 July 2014
   V2.0.2: Returns weights also when frac=0 for consistency.
       MC, Oxford, 3 November 2014
   V2.0.3: Updated documentation. Minor polishing. MC, Oxford, 8 December 2014
   V2.0.4: Fixed deprecation warning in Numpy 1.11. MC, Oxford, 18 April 2016
   V2.0.5: Fixed FutureWarning in Numpy 1.14. MC, Oxford, 18 January 2018
   V2.0.6: Dropped support for Python 2.7. MC, Oxford, 21 May 2018
   V2.1.0: Allow one to specify output coordinates different from the input ones.
       MC, Oxford, 20 July 2021
   Vx.x.x: additional changes are documented in the global package CHANGELOG.

"""

import numpy as np


################################################################################


class polyfit2d:

    def __init__(self, x, y, z, degree, weights):
        """
        Fit a bivariate polynomial of DEGREE 1 or 2 to a set of points
        (X, Y, Z) assuming errors in the Z variable only and weights=1/sigz^2.

        With DEGREE=1 this function fits a plane

           z = a + b*x + c*y

        while with DEGREE=2 the function fits a quadratic surface

           z = a + b*x + c*y + d*x*y + e*x^2 + f*y^2

        """
        a = np.column_stack([np.ones_like(x), x, y])
        if degree == 2:
            a = np.column_stack([a, x*y, x**2, y**2])

        sqw = np.sqrt(weights)
        self.degree = degree
        self.coeff = np.linalg.lstsq(a*sqw[:, None], z*sqw, rcond=None)[0]
        self.zfit = a @ self.coeff


    def eval(self, x, y):
        """ Evaluate at the point (x,y) the polynomial previously fitted """

        a = [1, x, y]
        if self.degree == 2:
            a = np.append(a, [x*y, x**2, y**2])

        zout = a @ self.coeff

        return zout


################################################################################


def biweight_sigma(y, zero=False):
    """
    Biweight estimate of the scale (standard deviation).
    Implements the approach described in
    "Understanding Robust and Exploratory Data Analysis"
    Hoaglin, Mosteller, Tukey ed., 1983, Chapter 12B, pg. 417

    """
    y = np.ravel(y)
    if zero:
        d = y
    else:
        d = y - np.median(y)

    mad = np.median(np.abs(d))
    u2 = (d / (9.*mad))**2  # c = 9
    good = u2 < 1.
    u1 = 1. - u2[good]
    num = y.size * ((d[good]*u1**2)**2).sum()
    den = (u1*(1. - 5.*u2[good])).sum()
    sigma = np.sqrt(num/(den*(den - 1.)))  # see note in above reference

    return sigma

################################################################################

def biweight_mean(y, itmax=10):
    """
    Biweight estimate of the location (mean).
    Implements the approach described in
    "Understanding Robust and Exploratory Data Analysis"
    Hoaglin, Mosteller, Tukey ed., 1983

    """
    y = np.ravel(y)
    c = 6.
    fracmin = 0.03*np.sqrt(0.5/(len(y) - 1.))
    y0 = np.median(y)
    mad = np.median(abs(y - y0))

    for it in range(itmax):
        u2 = ((y - y0)/(c*mad))**2
        u2 = u2.clip(0, 1)
        w = (1. - u2)**2
        w /= np.sum(w)
        mad_old = mad
        y0 += np.sum(w*(y - y0))
        mad = np.median(abs(y - y0))
        frac = abs(mad_old - mad)/mad
        if frac < fracmin:
            break

    return y0

################################################################################

def rotate_points(x, y, ang):
    """
    Rotates points conter-clockwise by an angle ANG in degrees.
    Michele cappellari, Paranal, 10 November 2013

    """
    theta = np.radians(ang)
    xNew = x*np.cos(theta) - y*np.sin(theta)
    yNew = x*np.sin(theta) + y*np.cos(theta)

    return xNew, yNew

################################################################################

def loess_2d(x, y, z, xnew=None, ynew=None,
             degree=1, frac=0.5, npoints=None, rescale=False, sigz=None):
    """
    loess_2d
    ========

    Purpose
    -------

    Two-dimensional LOESS smoothing via robust locally-weighted regression.

    This function is the implementation by `Cappellari et al. (2013)
    <https://ui.adsabs.harvard.edu/abs/2013MNRAS.432.1862C>`_ of the
    algorithm by `Cleveland (1979) <https://doi.org/10.2307/2286407>`_
    for the one-dimensional case and `Cleveland & Devlin (1988)
    <https://doi.org/10.2307/2289282>`_ for the two-dimensional case.

    Calling Sequence
    ----------------

    .. code-block:: python

       zout, wout = loess_2d(x, y, z, xnew=None, ynew=None, degree=1, frac=0.5,
                             npoints=None, rescale=False, sigz=None)

    Input Parameters
    ----------------

    x: array_like with shape (n,)
        vector of ``x`` coordinates.
    y: array_like with shape (n,)
        vector of ``y`` coordinates.
    z: array_like with shape (n,)
        vector of ``z`` coordinates to be LOESS smoothed.

    Optional Keywords
    -----------------

    xnew: array_like with shape (m,), optional
        Vector with the ``x`` coordinates at which to compute the smoothed
        ``z`` values.
    ynew: array_like with shape (m,), optional
        Vector with the ``y`` coordinates at which to compute the smoothed
        ``z`` values.
    degree: {1, 2}, optional
        degree of the local 2-dim polynomial approximation (default ``degree=1``).
    frac: float, optional
        Fraction of points to consider in the local approximation (default ``frac=0.5``).
        Typical values are between ``frac~0.2-0.8``. Note that the values are
        weighted by a Gaussian function of their distance from the point under
        consideration. This implies that the effective fraction of points
        contributing to a given value is much smaller that ``frac``.
    npoints: int, optional
        Number of points to consider in the local approximation.
        This is an alternative to using ``frac=npoints/x.size``.
    rescale: bool, optional
        Rotate the ``(x, y)`` coordinates to make the ``x`` axis the axis of
        maximum variance. Subsequently scale the coordinates to have equal
        variance along both axes. Then perform the local regressions.
        This is recommended when the distribution of points is elongated or
        when the units are very different for the two axes.
    sigz: array_like with shape (n,)
        1-sigma errors for the ``z`` values. If this keyword is used
        the biweight fit is done assuming these errors. If this keyword
        is *not* used, the biweight fit determines the errors in ``z``
        from the scatter of the neighbouring points.

    Output Parameters
    -----------------

    zout: array_like with shape (n,)
        Vector of smoothed ``z`` values at the coordinates ``(x, y)``, or at
        ``(xnew, ynew)`` if the latter are given as input. In the latter case
        ``zout`` has shape ``(m,)``.
    wout: array_like with shape (n,)
        Vector of biweights used in the local regressions. This can be used to
        identify outliers: ``wout=0`` for outliers with deviations ``>4sigma``.

        When passing as input the ``(xnew, ynew)`` coordinates, this output is
        meaningless and is arbitrarily set to unity.

    ###########################################################################
    """

    if frac == 0:
        return z, np.ones_like(z)

    assert x.size == y.size == z.size, 'Input vectors (x, y, z) must have the same size'

    if xnew is None:
        xnew, ynew = x, y

    assert xnew.size == ynew.size, 'Input vectors (xnew, ynew) must have the same size'

    if npoints is None:
        npoints = int(np.ceil(frac*x.size))

    if rescale:

        # Robust calculation of the axis of maximum variance
        #
        nsteps = 180
        angles = np.arange(nsteps)
        sig = np.zeros(nsteps)
        for j, ang in enumerate(angles):
            x_rot, y_rot = rotate_points(x, y, ang)
            sig[j] = biweight_sigma(x_rot)
        k = np.argmax(sig)  # Find index of max value

        x_rot, y_rot = rotate_points(x, y, angles[k])
        mx, my = biweight_mean(x_rot), biweight_mean(y_rot)
        sx, sy = biweight_sigma(x_rot), biweight_sigma(y_rot)
        x, y = (x_rot - mx)/sx, (y_rot - my)/sy

        xnew_rot, ynew_rot = rotate_points(xnew, ynew, angles[k])
        xnew, ynew = (xnew_rot - mx)/sx, (ynew_rot - my)/sy

    zout = np.empty_like(xnew, dtype=float)
    wout = np.empty_like(zout)

    for j, (xj, yj) in enumerate(zip(xnew, ynew)):

        dist = np.sqrt((x - xj)**2 + (y - yj)**2)
        w = np.argsort(dist)[:npoints]
        dist_weights = (1 - (dist[w]/dist[w[-1]])**3)**3  # tricube function distance weights
        zfit = polyfit2d(x[w], y[w], z[w], degree, dist_weights).zfit

        # Robust fit from Sec.2 of Cleveland (1979)
        # Use errors if those are known.
        #
        bad = None
        for p in range(10):  # do at most 10 iterations
        
            if sigz is None:                # Errors are unknown
                aerr = np.abs(zfit - z[w])  # Note ABS()
                mad = np.median(aerr)       # Characteristic scale
                uu = (aerr/(6*mad))**2      # For a Gaussian: sigma=1.4826*MAD
            else:                           # Errors are assumed known
                uu = ((zfit - z[w])/(4*sigz[w]))**2  # 4*sig ~ 6*mad
                
            uu = uu.clip(0, 1)
            biweights = (1 - uu)**2
            tot_weights = dist_weights*biweights
            poly = polyfit2d(x[w], y[w], z[w], degree, tot_weights)
            zfit = poly.zfit
            bad_old = bad
            bad = biweights < 0.34          # 99% confidence outliers
            if np.array_equal(bad_old, bad):
                break

        if np.array_equal(x, xnew):
            zout[j] = zfit[0]
            wout[j] = biweights[0]
        else:
            zout[j] = poly.eval(xj, yj)
            wout[j] = 1

    return zout, wout

################################################################################
```
